# Evolutionary priming and transition to the ectomycorrhizal habit in an iconic lineage of mushroom-forming fungi: is preadaptation a requirement?

**DOI:** 10.1101/2021.02.23.432530

**Authors:** Brian Looney, Shingo Miyauchi, Emmanuelle Morin, Elodie Drula, Pierre Emmanuel Courty, Annegret Kohler, Alan Kuo, Kurt LaButti, Jasmyn Pangilinan, Anna Lipzen, Robert Riley, William Andreopoulos, Guifen He, Jenifer Johnson, Matt Nolan, Andrew Tritt, Kerrie W. Barry, Igor V. Grigoriev, László G. Nagy, David Hibbett, Bernard Henrissat, P. Brandon Matheny, Jessy Labbé, Francis M. Martin

## Abstract

The ectomycorrhizal symbiosis is an essential guild of many forested ecosystems and has a dynamic evolutionary history across kingdom Fungi, having independently evolved from diverse types of saprotrophic ancestors. In this study, we seek to identify genomic features of the transition to the ectomycorrhizal habit within the Russulaceae, one of the most diverse lineages of ectomycorrhizal fungi. We present comparative analyses of the pangenome and gene repertoires of 21 species across the order Russulales, including a closely related saprotrophic member of Russulaceae. The ectomycorrhizal Russulaceae is inferred to have originated around the Cretaceous-Paleogene extinction event (73.6-60.1 million years ago (MY)). The genomes of the ectomycorrhizal Russulaceae are characterized by a loss of genes for plant cell-wall degrading enzymes (PCWDEs), an expansion of genome size through increased transposable element (TE) content, a reduction in secondary metabolism clusters, and an association of genes coding for certain secreted proteins with TE “nests”. The saprotrophic sister group of the ectomycorrhizal Russulaceae, *Gloeopeniophorella convolvens*, possesses some of these aspects (e.g., loss of some PCWDE and protease orthologs, TE expansion, reduction in secondary metabolism clusters), resulting from an accelerated rate of gene evolution in the shared ancestor of Russulaceae that predates the evolution of the ectomycorrhizal habit. Genomes of Russulaceae possess a high degree of synteny, including a conserved set of terpene secondary metabolite gene clusters. We hypothesize that the evolution of the ectomycorrhizal habit requires premodification of the genome for plant root association followed by an accelerated rate of gene evolution within the secretome for host-defense circumvention and symbiosis establishment.

## Introduction

Fungi fulfill diverse and essential functional roles in facilitating ecosystem viability at a multitude of scales, and these roles are directly mediated by their evolutionary history. Current understandings of functional roles of fungi are closely linked with their nutrition uptake mode because fungi must live in close proximity to nutrient sources for absorption. Fungal strategies for nutritent acquisition are dynamic across the fungal tree of life. Seemingly redundant trophic strategies have independently evolved numerous times. Within a single order, Sebacinales, we see multiple origins of plant-associated roles including endophytism, ectomycorrhizae, orchid mycorrhizae, ericoid mycorrhizae, and liverwort symbiosis all derived from a saprotrophic ancestry (Weiß et al. 2016). This plasticity of nutritional mode transition, though accentuated in Sebacinales, can be seen throughout the Agaricomycetes (Hibbett 2006). Molecular traits contributing to these plant-associated functional roles are largely unexplored, especially in ectomycorrhizal fungi (Koide et al. 2014; Martin et al. 2016).

The ectomycorrhizal (ECM) symbiosis is characterized by the transfer of water and nutrients to the plant and photoassimilates to the fungus through a cell-to-cell interface within roots, called the Hartig net (Smith & Read 2010). ECM has independently evolved in up to 82 lineages of fungi in Endogonomycetes, Pezizomycetes, and Agaricomycetes as well as 30 lineages of plants in Gymnospermae and Angiospermae (Tedersoo & Smith 2017; Brundrett & Tedersoo 2018). ECM fungi have evolved from diverse ancestral trophic states, including white-rot saprotrophs, brown-rot saprotrophs, litter decomposers, and root endophytes, with each evolutionary history necessitating different selective pressures (Tedersoo & Smith 2013; Martin et al. 2016; Pellitier & Zak 2018; Strullu-Derrien et al. 2018; Miyauchi et al. 2020). These evolutionary shifts in trophic strategy often led to specializations of function that contribute to changes in diversification rates that are defining for clades of fungi (Sánchez-García et al. 2017; 2020; Lutzoni et al. 2018).

The characterization of the *Laccaria bicolor* genome established a number of attributes for the genome of an ECM fungus, such as a high TE content, loss of plant cell-wall degrading enzymes (PCWDEs), and occurence of effector-like mycorrhiza-induced small secreted proteins (MiSSPs) during symbiosis (Martin et al. 2008; Labbé et al. 2012; Pellegrin et al. 2015; Plett et al. 2017). These genomic features have since been found in a wide array of mycorrhizal fungi belonging to Ascomycota, Basidiomycota and Mucoromycota (Kohler et al. 2015; Martin et al. 2016; Morin et al. 2019; Miyauchi et al. 2020). To more precisely establish the evolutionary events defining the origin(s) of ECM associations and to discriminate these from lineage-specific evolutionary changes, comparative genomic analyses of densely sampled evolutionary lineages of ECM fungi, all descended from a single origin of symbiosis, are needed. The evolution of ECM genomes within a single, densely sampled lineage has thus far only been investigated for the Amanitaceae, which showed a rapid expansion and contraction of functionally relevant genes early in the evolution of the ECM habit (Hess et al. 2018).

Russulaceae is an iconic lineage of ECM fungi that are dominant in ectotrophic landscapes and are prized for their edible mushrooms (Looney et al. 2018). Russulaceae possesses several ecologically relevant attributes that warrant study in a genomic context, such as a nitrophilic tendency of some members (Lilleskov et al. 2002), the production of unique sesquiterpenoid secondary compounds (Clericuzio et al. 2012), and an accelerated evolutionary rate of speciation, morphological transition, and host expansion (Looney et al. 2016). The vast majority of Russulaceae are ECM and mushroom-forming. However, the sister group of the ECM clade is a group of wood-decaying crust fungi of which we have sampled *Gloeopeniophorella convolvens* (Larsson & Larsson 2003). Thus, Russulaceae provides an exceptional opportunity to study an evolutionary transformation between nutritional modes and fruiting body forms. Here, we describe trends in genomic architecture and gene content in twenty representative species of Russulales. Our dataset contains fourteen previously unanalyzed genomes, including 11 species of ECM Russulaceae and their saprotrophic sister group (*Gloeopeniophorella convolvens*), and *Amylostereum chailletii*, a white-rot wood-decomposer that is associated with siricid woodwasps in a timber pathogenic symbiosis (Fitza et al. 2016). Our analysis elucidates patterns of functional diversity that have evolved within the ECM symbiotrophs, including evolution of PCWDEs, retention of genes to scavenge nitrogen compounds in soil organic matter, secondary metabolism, and TE invasion favoring duplication of species-specific genes. We hypothesize that a defined core set of genes derived from the common ancestor of ECM Russulaceae defines a particular niche for this lineage according to the ‘family gene conservation’ hypothesis (Looney et al. 2018).

## Results

### Phylogeny of Russulales

A reconstructed phylogeny of Russulales fungi showed members of *Lactarius* formed the sister clade to the rest of ECM Russulaceae, and members of *Lactifluus* formed the sister clade to a clade comprised of *Russula* (Fig. 1a, Fig. S1). Within *Russula*, a clade comprised of *R. brevipes* and *R. dissimulans* was inferred as sister to the rest of *Russula* (Fig. S1). The common ancestor of the ECM habit in Russulaceae is inferred to have arisen around the Cretaceous-Paleogene (K-Pg) extinction event (73.6-60.1 MY), a period of rapid ecological and anatomical innovation in plant communities (Alfaro et al. 2018). The family Russulaceae, including the saprotrophic *Gloeopeniophorella convolvens*, began diversification around the same time as the saprotrophic Auriscalpiaceae during the Cretaceous (~74 MY).

**Figure 1.**
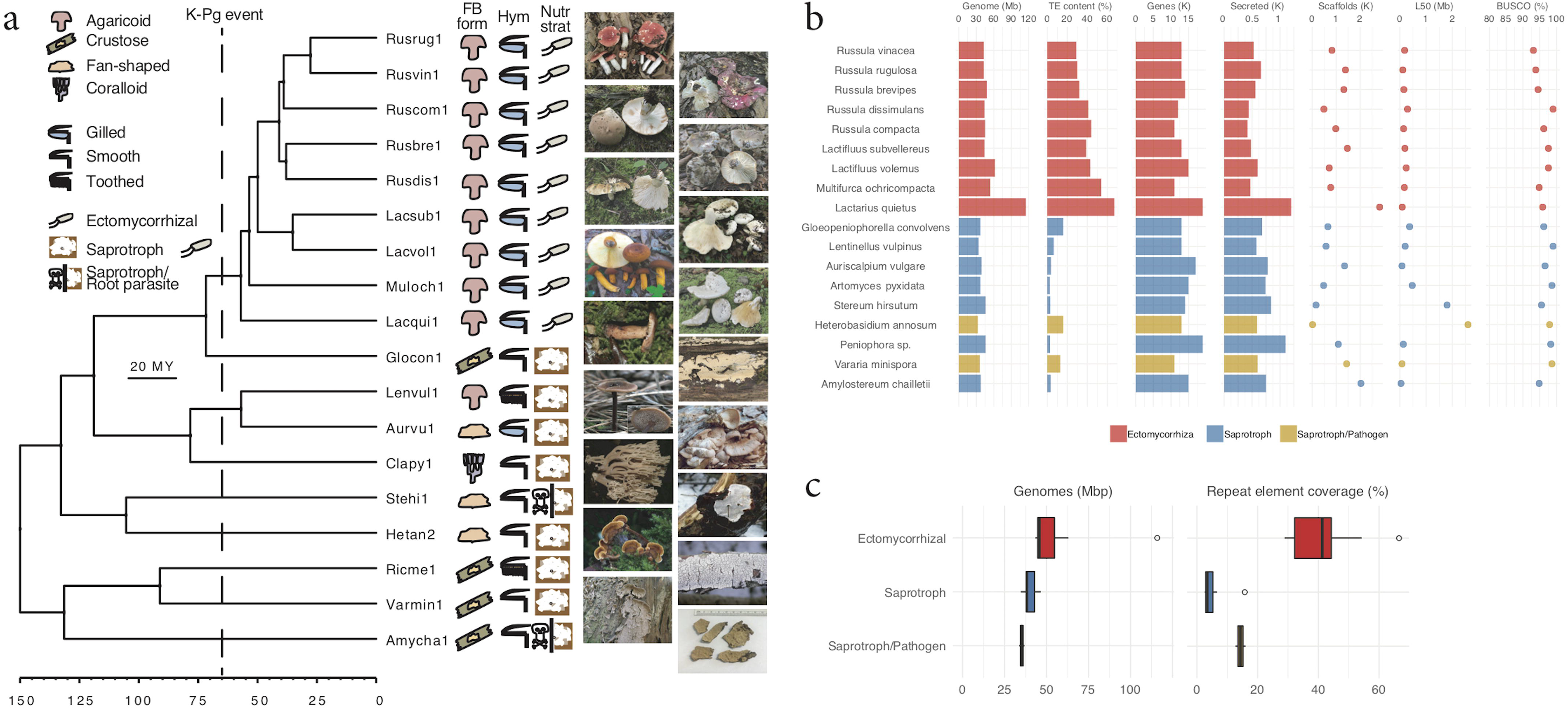
Overview of genome features of 18 fungi. a. Phylogenetic reconstruction of 18 Russulales genomes. 2,518 single-copy genes were used in RAxML with 1,000 bootstrap iterations. Taxon labels correspond to JGI identifiers Rusvin (*R. vinacea*), Rusrug (*R. rugulosa*), Rusbre (*R. brevipes*), Rusdis (*R. dissimulans*), Ruscom (*R. compacta*), Lacsub (*L. subvellereus*), Lacvol (*L. volemus*), Muloch (*M. ochricompacta*), Lacqui (*L. quietus*), Glocon (*G. convolvens*), Lenvul (*L. vulpinus*), Aurvu (*A. vulgare*), Clapy (*A. pyxidata*), Stehi (*S. hirsutum*), Hetan (*H. annosum*), Ricme (*Peniophora sp.*), Varmin (*V. minispora*), Amycha (*A. chailletii*). Fruitbody form (FB form), hymenium type (Hym), and nutritional strategy (Nutr strat) are given with images of the genome source. See details (Fig. S1) b. Genome: Genome size. TE content: coverage of transposable elements in the genomes. Genes: number of genes. Secreted: number of predicted secreted proteins (see Methods). Scaffolds: number of scaffolds. L50: N50 length. BUSCO: Genome completeness (Table S7). c. Genome size with repeat element coverage per ecological group.

### Main genomic features of Russulales

Genomes of ECM species within Russulales are larger than saprotrophic/pathogenic species (p<0.01, Wilcoxon signed-rank (WSR) test, 2-tailed), with *Lactarius quietus* having the largest genome (115.9 Mb) and other ECM genomes ranging from 43.3 to 90.3 Mb (Fig. 1b; Fig. 1c; Table S7). Over 94% of a benchmark set of conserved fungal genes (BUSCO, Simão et al. 2015) were found in genome assemblies (Fig. 1b), and up to 97% of the RNA-Seq reads mapped to the gene repertoire (see Info page at JGI Russulales portal, https://mycocosm.jgi.doe.gov/Russulales/Russulales.info.html), indicating that assembled genomes capture most of the coding gene space.

The coding pangenome of Russulales comprises over 250,000 predicted genes for the eighteen species compared, ranging from 10,514 genes for *Multifurca ochricompacta* to 18,952 genes for *Peniophora* sp. (Fig. S3). The common, conserved genes, which are shared among the eighteen fungi, including some missing from one or fewer species, make up one quarter of all genes ranging around 3,500 genes for most species and up to 4,023 genes for *L. quietus*. The species-specific gene content varies considerably between species but not trophic categories, with *Peniophora* sp. and *L. quietus* having the highest number of unique genes (11,721 and 10,313 respectively), and *M. ochricompacta* with only 2,832 unique genes. Secondary alleles were identified from sequenced dikaryotic genomes by the PacBio sequencing technology; they comprised from 14 to 39% of all protein models (Table S7).

### The evolution of plant cell wall degrading enzymes in ECM Russulales

Copy number of gene families in the predicted secretome likely reflects evolutionary adaptations (Martin et al. 2016). We annotated and manually curated the whole set of genes coding for carbohydrate-active enzymes (CAZymes), using the CAZy database (Lombard et al. 2014). ECM Russulales contain a smaller set of CAZymes than saprotrophic taxa (Fig. 2; see Fig. S4 for Individual CAZymes). They have lost a core set of genes required for efficient degradation of PCWDEs and fungal cell wall degrading enzymes (FCWDEs). The number of gene copies for many secreted enzymes involved in the decomposition of cellulose, hemicellulose, pectin, lignin, chitin and mannan is restricted or absent in symbiotrophs compared to the taxonomically related saprotrophs, including *G. convolvens* (Table 1; Fig. S4; Table S16). For many orthologous clusters, however, this reduction is seen to occur in the ancestor of Russulaceae, including *G. convolvens* (Table S16). These orthogroups include subtilases, aspartic proteases, AA3_2 aryl alcohol oxidases, GH12 endoglucanases, and expansin-like proteins among others. The ECM Russulales have kept a few orthologous clusters involved in cellulose degradation, such as glycoside hydrolases GH45 and lytic polysaccharide monooxygenases (LPMOs, AA9) that may be involved in the host root penetration or fungal cell wall remodeling (Table S17)(Veneault-Fourrey et al. 2014; Krizsán et al. 2019; Zhang et al. 2019).

**Figure 2.**
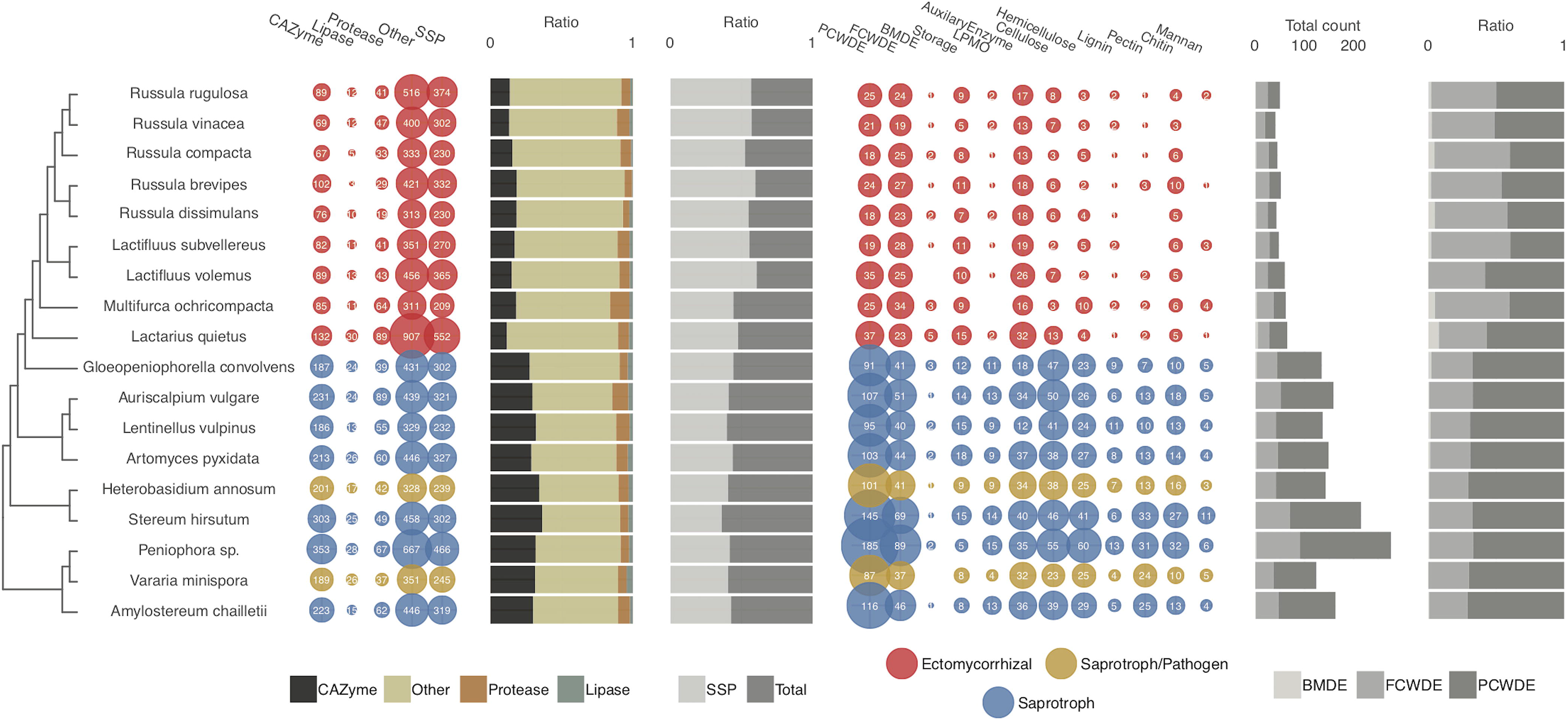
Predicted secretomes of 18 members of Russulales. First bubble plot (left): The number of secreted genes for CAZymes, lipases, proteases, and others (i.e., all secreted proteins not in these first three groups). The group SSPs is a subcategory showing the number of small secreted proteins (< 300 aa). The size of bubbles corresponds to the number of genes. The fungi are coloured according to their ecology. First bar plots (middle): The ratio of CAZymes, lipases, proteases, to all secreted proteins (left); and the ratio of SSPs among the entire secretome (right). Second bubble plot (right): The number of including plant cell wall degrading enzymes (PCWDE) and microbial cell wall degrading enzymes (MCWDE), bacterial membrane (i.e., peptidoglycan) degrading enzyme (BMDE), lytic polysaccharide monooxygenase (LPMO), enzymes for starch and glycogen (storage); AA family CAZymes (Auxiliary Enzymes); substrate-specific enzymes for cellulose, hemicellulose, lignin, and pectin (plant cell walls); chitin, glucan, mannan (fungal cell walls). Second bar plots (far right): The total count of genes including PCWDE, MCWDE, and BMDE (left); and the proportion of PCWDE, MCWDE, and BMDE (right) (Table S4; Table S9).

**Table 1.** Total and secreted gene families enriched for species of ECM Russulaceae and functional groups.

|  | Total gene repertoire | Secreted proteins |
| --- | --- | --- |
| Lacpsa | **GH13_32** (α-amylase), **GH15** (glucoamylase), **GH152** (glucanase), **GT3** (glycogen synthase), **A02A** (aspartyl protease), **C97** (desumoylating isopeptidase), **I29** (cathepsin propeptide), **I87** (peptidase), **M17** (leucyl aminopeptidase), **M36** (fungalsin), **M43B** (metallo-endopeptidase), **I13** (serine protease inhibitor), **S11** (serine protease), **A08**, **AA14**, **A28A**, **C110**, **C111**, **C115**, **C11X**, **C82A**, **C83**, **C95**, **I15**, **I16**, **I17**, **I71**, **I83**, **M15B**, **M20F**, **M28F**, **M42**, **M48C**, **M50A**, **M82**, **M87**, **M96**, **M98**, **P01**, **P02A**, **P02B**, **S49B**, **S49C**, **S81**, **U32**, **U62**, **U74** | **CBM50** (carbohydrate-binding module), **EXPAN** (expansin), **GH15** (glucoamylase), **M36** (fungalsin), **M43B** (metallo-endopeptidase), **GH152**, **M28F**, **S09B**, **S41A** |
|  | AA14, CBM12, CBM20, CE4, EXPN, GH13_32, GH15, GH152, GT3, PL35, A02A, C04, C45, C67, C97, I01, I04, I08, I09, I25B, I29, I31, I32, I43, I51, I87, M16B, M17, M36, M43B, S09B, I13, S41A, C89, S11, A08, A28A, C110, C111, C115, C11X, C40, C82A, C83, C95, I15, I16, I17, I71, I83, M15B, M20F, M28F, M42, M48C, M50A, M82, M87, M96, M98, P01, P02A, P02B, S49B, S49C, S81, U32, U62, U74, GGGX\|ABH03 | AA3_2, AA14, CBM50, EXPN, GH15, GH152, S10, C19, I08, M28F, S09B, S41A, GGGX\|ABH03 |
| Lacqui | **AA3_3** (alcohol oxidase), **AA5_1** (copper radical oxidase), **CBM13** (carbohydrate-binding module), **GH45** (cellulase), **GH92** (mannosidase), **GT20** (α-trehalose-phosphate synthase), **GT22** (mannosyltransferase), **GT24** (glycoprotein α-glucosyltransferase), **C54** (cysteine protease), **S53** (sedolisin), **T06** (threonine protease) | **AA5_1** (copper radical oxidase), **EXPAN** (expansin), **GH45** (cellulase), **S53** (sedolisin), **GH13**, **GH13_22**, **I63**, **M24A**, **T06** |
|  | AA1, AA3, AA3_3, AA5_1, CBM12, CBM13, EXPN, GH13, GH13_22, GH25, GH45, GH92, GT20, GT22, GT24, GT76, C04, C54, C65, M14A, M16C, S53, T06, GGGX\|ABH03 | AA3_2, AA5_1, EXPN, GH13, GH13_22, GH25, GH45, S10, S53, I02, I63, M24A, T06, GGGX\|ABH03 |
| Lacsub | **AA5_1** (copper radical oxidase), **GH37** (trehalase) | **I25B** |
|  | AA5_1, CBM12, GH13_22, GH37, M13, S09C, GX\|ABH08 | AA5_1, S09C, M23B, I25B, I25X |
| Lacvol | **AA1_1** (laccase), **GH20** (β-N-acetylglucosaminidase), **GH37** (trehalase), **S28** (lysosomal Pro-Xaa carboxypeptidase) | **AA1_1** (laccase) |
|  | AA1_1, AA1_2, GH13_22, GH15, GH20, GH37, GH38, | AA1_1, GH9, S10, |
|  | GT66, A22B, C46, M03A, M76, S28, S72, T02, T03, T06, C82, GGGX\|ABH03 | GGGX\|ABH03 |
| Muloch | **S08A** (subtilisin), **S28** (lysosomal Pro-Xaa carboxypeptidase) | **CE4** (chitin deacetylase), **GH47** (α-mannosidase), **M43B** (cytophagalyisin), **S08A** (subtilisin), **CE14** |
|  | CE4, CE14, GH47, GT22, C01B, I25A, M43B, S08A, S28, GX\|ABH07 | CE4, CE14, GH47, M43B, S08A, I25X |
| Rusbre | **CBM50** (carbohydrate-binding module), **GH47** (α-mannosidase), **I43** (serine protease inhibitor), **AA5** | **CBM50** (carbohydrate-binding module) |
|  | AA5, CBM18, CBM43, CBM50, CE14, GH3, GH5_30, GH17, GH47, GH72, GT39, M23B, C82, A31, I32, I43 | AA1_1, CBM50, GH9 |
| Ruscom |  |  |
|  | GH13_1, GH13_5, A31 | AA5_1, CBM50 |
| Rusdis |  |  |
|  | AA1_2, CE9, GH13_1, C39, I32, M48X, S09A | GH13_32, I25X |
| Rusrug | **GH5_30** (cellulase), **GT48** (1,3-β-glucan synthase), **C19** (ubiquitin-specific protease), **S16** (Ion protease), **C85** | **GH5_12** (cellulase), **GH5_30** (cellulase) |
|  | CBM12, EXPN, GH5_30, GH30, GH92, GT31, GT48, GT49, GT59, C19, C45, I32, S10, S16, C40, C85 | EXPN, GH5_12, GH5_30, S10, C19 |
| Rusvin | **GT4** (glycosyltransferase), **GT48** (glycosyltransferase), **C19** (ubiquitin-specific protease), **I21** | **GH5_30** (cellulase), **CE9** |
|  | AA7, CE9, EXPN, GT4, GT48, GT69, GT76, C19, I25X, I32, C89, A31, I21 | EXPN, CE9, GH5_30 |
| ECM Russulaceae | **GH45** (cellulase), **I32** (IAP) | **GH45** (cellulase) |
| Saprotrophs | **AA2**, **AA3_1**, **AA3_2**, **AA3_4**, **AA8**, **AA9**, **CBM1**, **CBM5**, **CBM35**, **CE1**, **CE8**, **CE15**, **CE16**, **GH1**, **GH2**, **GH3**, **GH5**, **GH5_5**, **GH5_7**, **GH5_12**, **GH5_22**, **GH5_50**, **GH6**, **GH7**, **GH10**, **GH11**, **GH12**, **GH16**, **GH18**, **GH27**, **GH28**, **GH29**, **GH30_3**, **GH31**, **GH43**, **GH51**, **GH53**, **GH55**, **GH74**, **GH76**, **GH78**, **GH79**, **GH81**, **GH95**, **GH105**, **GH115**, **GH128**, **GH131**, **GH145**, **GT41**, **PL8_4**, **PL14_4**, **CO3B**, **C12**, **C56**, **I51**, **M28E**, **G01**, **M77**, **GGGX\|ABH04**, **GX\|ABH09**, **GX\|ABH23** | **AA2**, **AA3_2**, **AA8**, **AA9**, **CBM1**, **CBM5**, **CBM35**, **CE1**, **CE8**, **CE15**, **CE16**, **GH3**, **GH5_5**, **GH5_7**, **GH6**, **GH7**, **GH10**, **GH12**, **GH18**, **GH28**, **GH30_3**, **GH35**, **GH43**, **GH51**, **GH53**, **GH55**, **GH72**, **GH79**, **GH81**, **GH92**, **GH95**, **GH115**, **GH145**, **PL8_4**, **PL14_4**, **G01** |

\* red indicates substantial enrichment; purple indicates a unique family; black indicates no depletion

Although ECM Russulales have experienced a concerted loss of CAZymes, members of Russulaceae have experienced species-specific expansions of particular gene families and orthologues (Table 1; Table S17). This includes enzymes involved in degradation of cellulose (e.g., GH5_12, GH5_30, GH45), hemicellulose (e.g., CE4, CBM13), chitin (e.g., GH20, CBM18), and mannan (e.g., GH92). The second largest genome, *L. volemus*, possesses the highest copy numbers of AA1_1 laccase in Russulales. The third largest ECM Russulaceae genome, *M. ochricompacta*, possesses the fewest number of genes in Russulales and the second highest TE proportion. *Multifurcata ochricompacta* is particularly expanded in three groups of subtilisin-like serine proteases which are absent for most of the rest of Russulaceae and has also seen expansions of secreted CE4 chitin deacetylases and GH47 α-mannosidases. *Lactifluus subvellereus* is characterized by a substantial expansion of AA5_1 glyoxal oxidases with moderate expansions in aspartyl proteases and AA1_1 laccases. *Russula brevipes* is the only ECM Russulaceae species to possess PL14_4 β-1,4-glucuronan lyases and is expanded in GH72-CBM43 β-1,3-glucanosyltransglycosylases. Expansion in *R. rugulosa* includes a group of serine carboxypeptidases and a group of tyrosinases. *Russula vinacea* is highly expanded in a cluster of carboxylesterase lipases. Not all ECM Russulaceae species exhibit substantial expansions in their secretome, with *R. compacta* and *R. dissimulans* mostly lacking gene duplication-mediated expansions. Substantial expansions are less common for non-ECM Russulales, with the exception of *Peniophora*, that sees substantial expansion in a cluster of lipases and AA1_1 laccases, and *A. vulgare* with expansions of two subtilase clusters, a GMC oxidoreductase cluster, and a cluster of aspartyl proteases.

The total number of secreted proteases in ECM Russulaceae was also significantly reduced compared to the saprotrophs, however symbiotrophs show expansions of aspartyl proteases, subtilisin-like serine proteases, and serine carboxypeptidases (Table 1; Table S17). The most extreme gene expansion of 283 genes in a single orthologous cluster of subtilases with pro-kumamolisin activation domains was observed for *L. quietus*. *Lactarius psammicola* possesses the most diverse repertoire of expanded and unique proteases including fungalysin, aspartyl proteases, and metallo-endopeptidases.

Functional specialization is also evident in unique species-specific secreted gene clusters for ECM Russulaceae that are highly enriched. *Lactarius quietus* possesses unique secreted gene clusters, with two clusters of putative fungistatic metabolites as well as a thaumatin-like protein group and a group of unique expansin-like proteins. *Multifurca ochricompacta* possesses a unique cluster of fungistatic metabolite genes. *Lactifluus volemus* possesses a unique cluster of secreted protein genes with a LysM domain. *Lactifluus subvellereus* possesses a unique cluster of GH45 endoglucanases. *Russula compacta* possesses a unique cluster of hydrophobin genes. *Russula brevipes* is characterized with seven unique gene clusters, of which the largest is a cluster of serine carboxypeptidases. *Russula dissimulans* possesses a cluster of glutathione S-transferases. *Russula rugulosa* is another genome with many expanded unique gene clusters, with a single large cluster of expansin-like proteins. *Peniophora* is the only non-ECM member with a large quantity of unique gene clusters, with 26 unique gene clusters of which the largest cluster has 9 gene copies.

At a gene family level, the ECM Russulaceae is enriched in GH45 cellulases and inhibitors of caspases and cysteine endopeptidases (family I32) compared with saprotrophic Russulales (Table 1). An important aspect of ECM Russulaceae specialization is the retention of lignolytic manganese peroxidase (POD) genes as a remnant of Russulaceae’s white rot ancestry (Fig. S16). These species have not retained the same POD genes, however, with two clades of POD genes having been recovered. The genes are split between both members of *Russula* and *Lactifluus*, indicating that there were multiple independent losses of these genes in both clades. In both cases, the same gene has been retained in the closest extant saprotrophic ancestor, *G. convolvens* making it less likely that they are functionally redundant due to recent gene duplication.

### The repertoire of small secreted proteins

The number of small secreted proteins (SSPs) with unknown function is similar in symbiotrophs and saprotrophs (see ratio of SSP and Total in Fig. 2). The secretome of *L. quietus* encodes 552 gene coding for SSPs, almost twice as high compared to other ECM Russulaceae, suggesting the possibility of a whole-genome duplication event or a drastic shift in functional specialization. We identified a SSP gene that is evolutionarily conserved in all ECM Russulaceae species. It encodes a phosphatidylglycerol/phosphatidylinositol transfer protein with an ML domain implicated in the detection of lipids and pathogenicity factors (Ph.tr.pr in red; Table S2).

### Secondary metabolite diversity

409 gene clusters involved in secondary metabolism were detected from 21 Russulales genomes (including the addition of the recently sequenced *L. psammicola* BPL869 v1.0, *L. indigo* 2018DUKE089 v1.0, and *R. earlei* BPL698 v1.0) (Fig. 3). Saprotrophic members of Russulales possess more NRPS-like (p<0.01, WSR test, 2-tailed), combined T1PKS & NRPS-like (p<0.001, WSR test, 2-tailed), and total number of clusters (p<0.01, WSR test, 2-tailed) than ECM species, whereas ECM members possess more siderophore clusters (p=0.03, WSR test, 2-tailed). Four NRPS clusters share homology among some or most of Russulaceae, which represent two different metabolic pathways (Fig. S17a). A diverse group of up to eight TPC clusters for terpene synthase are conserved among at least some members of Russulaceae, with two clusters also represented in Auriscalpiaceae (Fig. S17b). Two clusters representing a single PKSI pathway with a ferric reductase transporter domain as well as another group of two clusters representing a single pathway with an aspartyl protease domain is present for most members of Russulaceae, including the saprotrophic *G. convolvens* (Fig. S17c & S17d).

**Figure 3.**
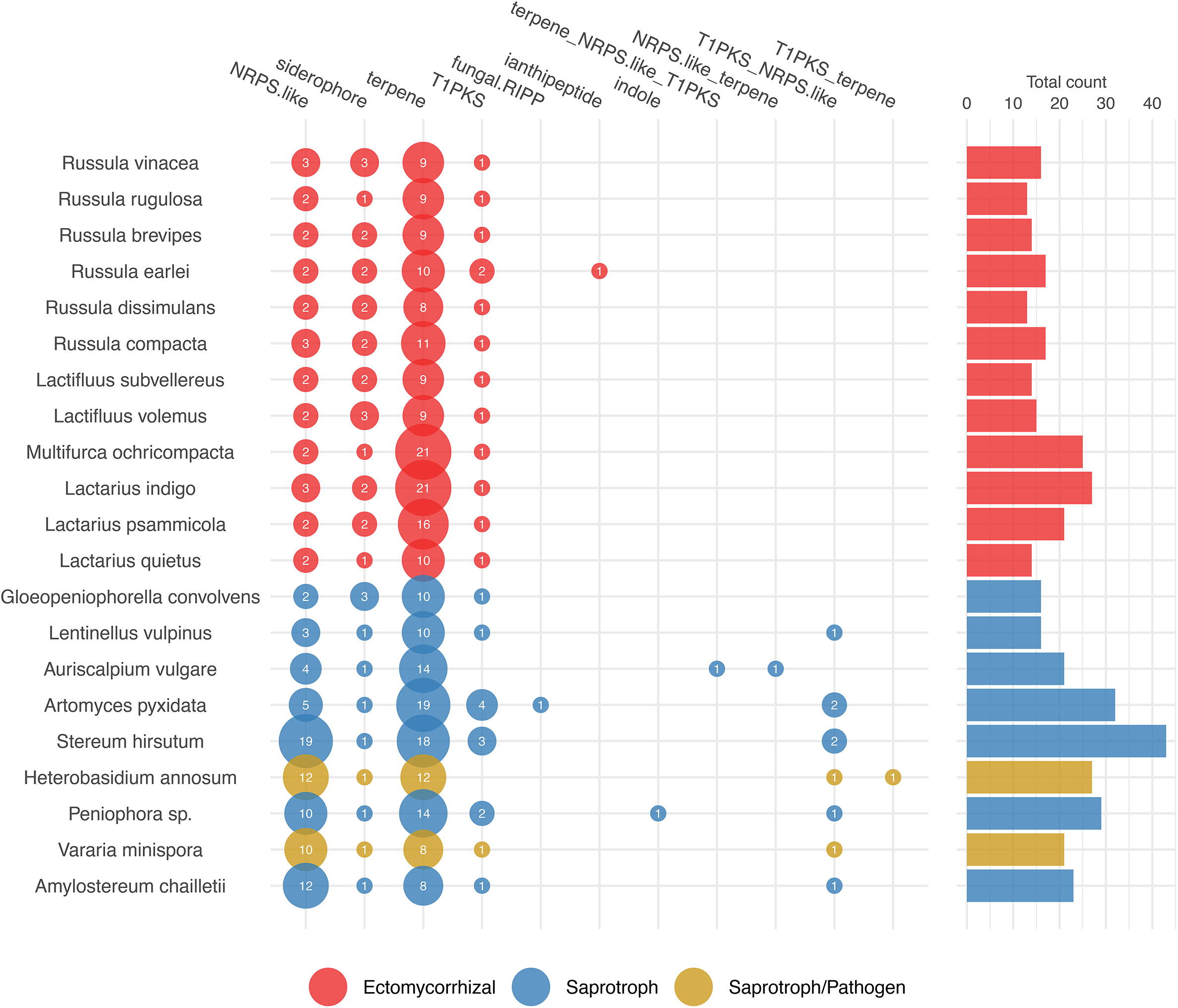
Predicted secondary metabolite biosynthesis gene clusters of 20 members of Russulales. a. The number of SMCs predicted for NRPS-like, siderophore, terpene, T1PKS, fungal-RIPP, ianthipeptide, indole, and hybrid classes (i.e., containing features of multiple classes). The size of the bubbles corresponds to the number of clusters. The fungi are coloured according to their ecology.

### Gene evolution rate in Russulaceae

Across the pangenome, ECM Russulaceae and saprotrophic Russulales experienced comparable gene duplications and contractions with a slightly lower overall rate of gene gain (Fig. 4). The overall rate of gene evolution was accelerated in the ancestors of both ECM Russulaceae (internode 8-9) and saprotrophic Russulaceae (internode 9-12). Species-specific gene evolutionary rates were higher for *L. quietus* and *S. hirsutum* across the pangenome and only for ECM Russulaceae members across the secretome. Gene evolution rate varied across the secretome, with a higher rate of gene loss (0.08), gene duplication rates at about half of the loss rate (0.04), and gene gain rates ten times less (0.004). An accelerated rate of gene loss occurred in the pangenome of the ancestor of Russulaceae (node 8).

**Figure 4.**
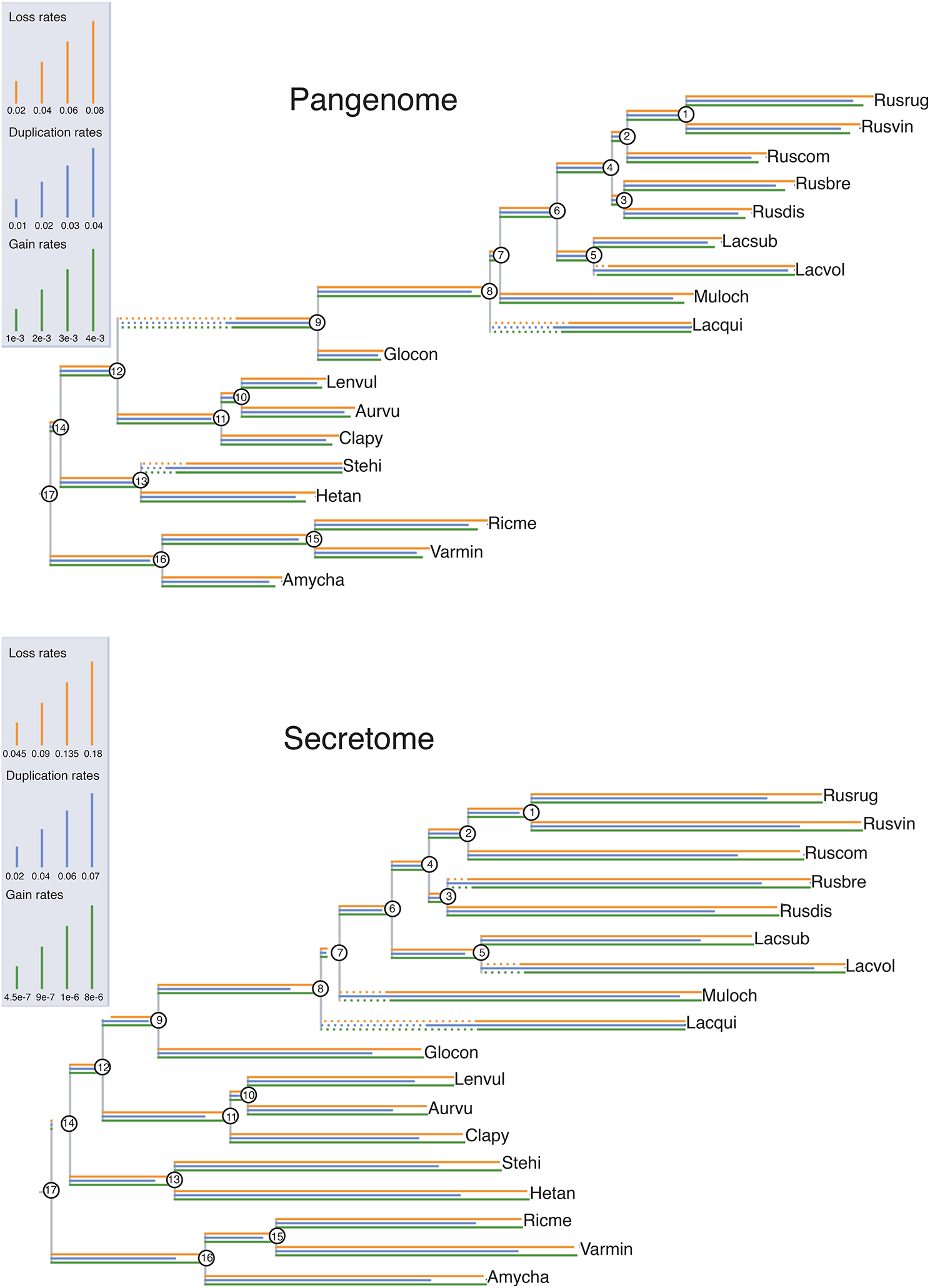
Evolutionary rate COUNT analysis of Russulales genomes. Top) Rates of gene loss, duplication, and gain for pangenomes along branches. Poisson distribution of birth-death model is given for the root node. Bottom) Rates of gene gain, loss, and duplication for the secretome along branches with dotted lines representing the rate length exceeding the total line. Ancestral nodes are numbered. The labels show short JGI fungal IDs. Rusvin (*R. vinacea*), Rusrug (*R. rugulosa*), Rusbre (*R. brevipes*), Rusdis (*R. dissimulans*), Ruscom (*R. compacta*), Lacsub (*L. subvellereus*), Lacvol (*L. volemus*), Muloch (*M. ochricompacta*), Lacpsa (*L. psammicola*), Lacqui (*L. quietus*), Glocon (*G. convolvens*), Lenvul (*L. vulpinus*), Aurvu (*A. vulgare*), Clapy (*A. pyxidata*), Stehi (*S. hirsutum*), Hetan (*H. annosum*), Ricme (*Peniophora* sp.), Varmin (*V. minispora*), Amycha (*A. chailletii*).

However, gene loss was the greatest in species-specific lineages, indicating a high evolutionary rate of secretome modification. Gene loss rates are over twice as high as the overall genome rates (0.18) and gene gain rates are three orders of magnitude lower than the overall genomes rates (8×10^−6^).

### Gene synteny in Russulales

Gene synteny of five *Russula* species was compared (Fig. 5). Russulaceae share the highest level of synteny with each other despite the fact that the size of scaffolds for the ten largest scaffolds is variable within Russulales genomes (Fig. S9-10). Syntenic regions are disrupted by clusters of TEs (Fig. 6). The frequency of TE insertions suggests that TEs accumulated further in TE-rich regions during the course of evolution, forming ‘transposon nests.’ The single largest syntenic block among five *Russula* species was used as a landmark to investigate gene order and mesosynteny (Fig. S14-15). We found that locations of TEs and unclassified repeats are aligned with SSPs in the syntenic regions. To examine the significance of this potential association, we performed a permutation test to compare distances between TEs and SSP coding genes of 18 Russulales. We found that TEs and SSPs of unknown function are significantly closer in ECM Russulaceae genomes than in non-ECM Russulales (p < 0.01; Fig. 7).

**Figure 5.**
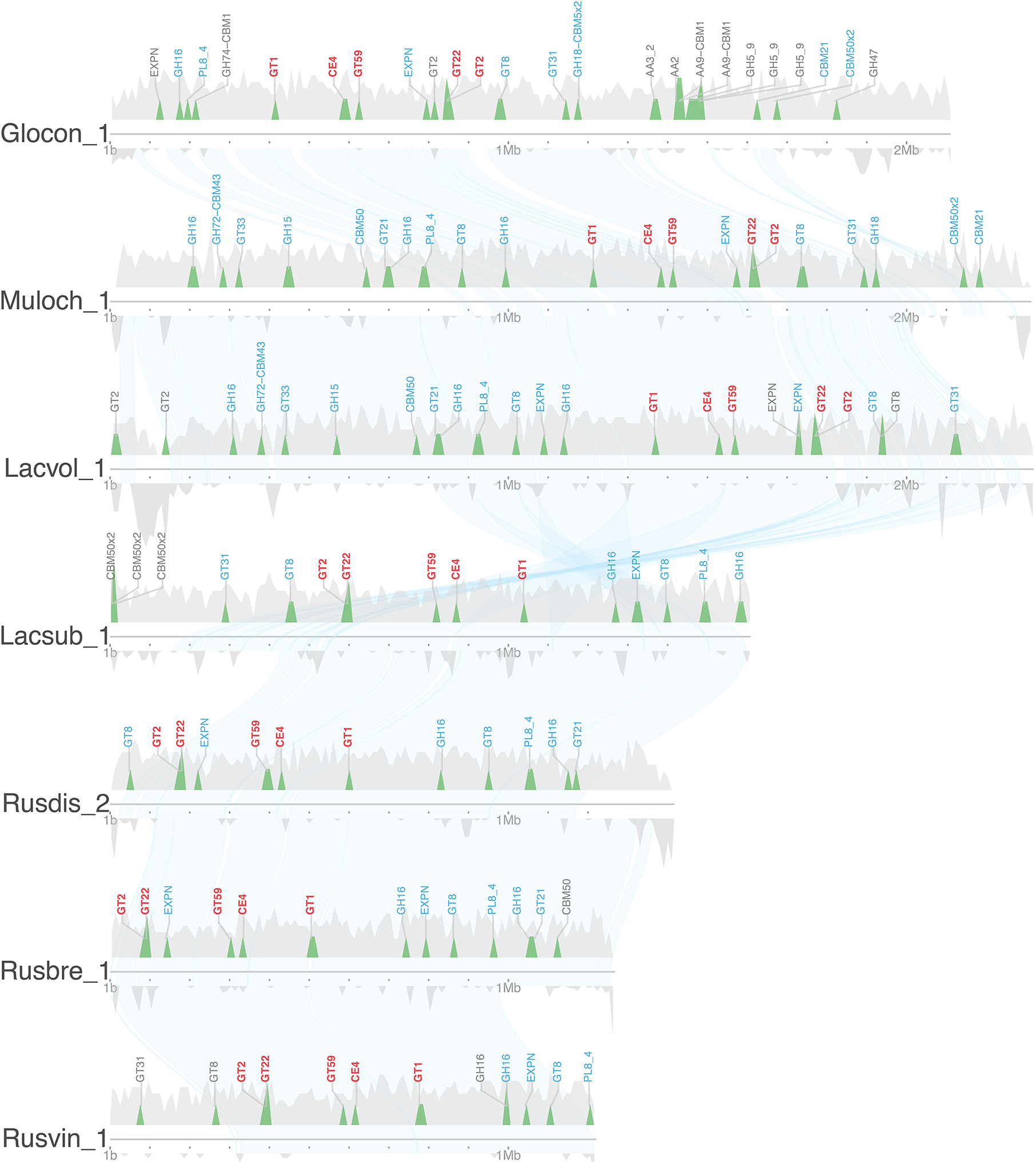
Genomic locations of genes for small, secreted proteins in syntenic regions. Scaffold 1 of *G. convolvens* (Glocon1) is aligned with other closely related fungi. Kirisame (drizzle) plot represents genes, secretome (genes for small, secreted proteins, CAZymes, proteases, lipases), and repeat elements in syntenic regions. Small, secreted proteins (SSP) coding genes are labelled. Upward peaks: Density of all genes (light grey) coded and genes coding for secreted proteins (colors). Downward peaks: Density of repeat elements including TEs and unknown repeats. Blue lines: Syntenic regions. Species are in evolutionary order. Scaffold ID: short JGI fungal ID with scaffold number. *G. convolvens* (Glocon), *M. ochricompacta* (Muloch), *L. volemus* (Lacvol), *L. subvellereus* (Lacsub), *R. dissimulans* (Rusdis), *R. brevipes* (Rusbre), *R. vinacea* (Rusvin).

**Figure 6.**
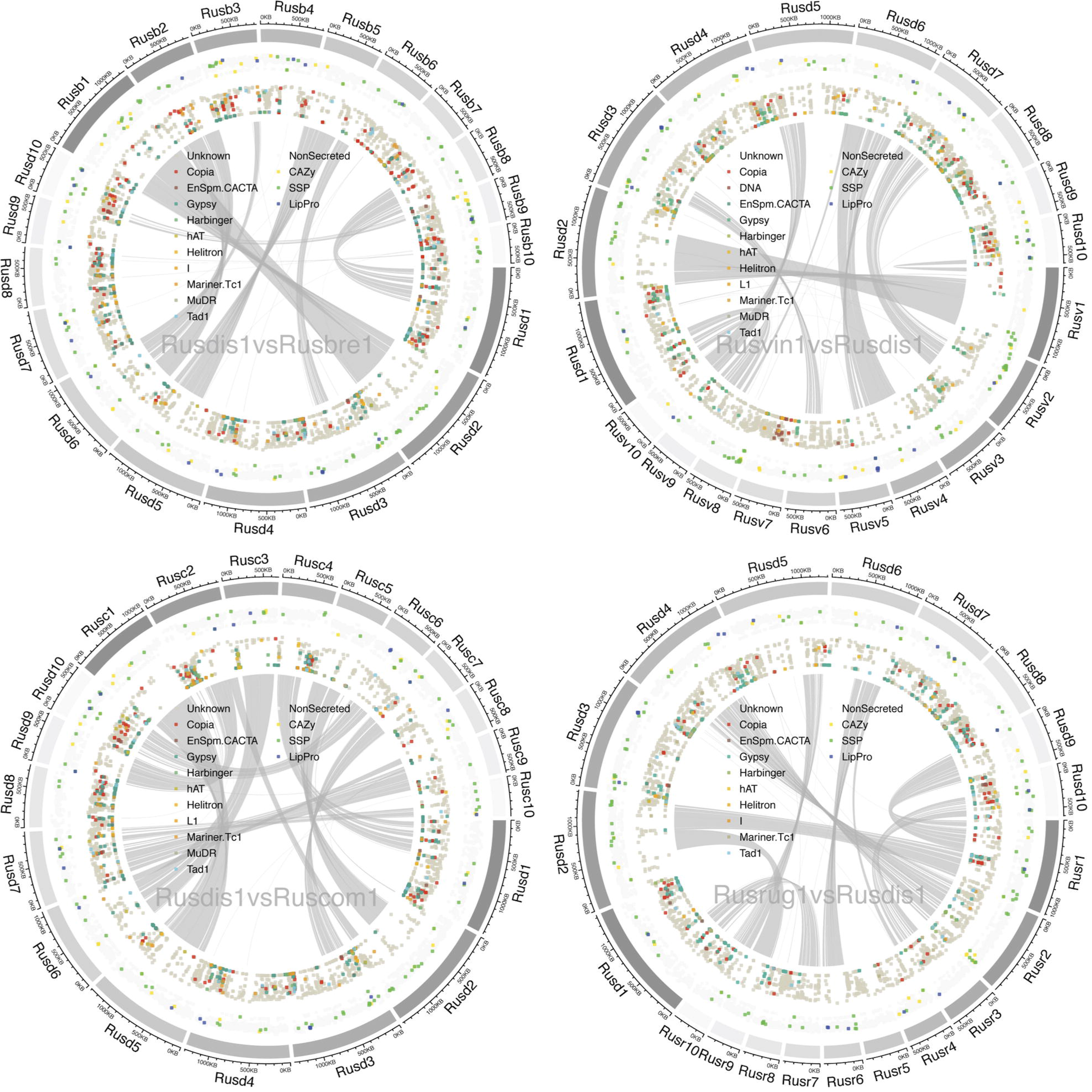
Macrosynteny comparison with five *Russula* species. Hanabi (firework) plot shows pairwise syntenic comparison of Scaffold 1 to 10. Outer circle: The size of scaffold 1 to 10. First inner circle: Genes located in the scaffolds. Genes coding for CAZymes, SSPs, lipases, proteases are highlighted (see the legend for details). Second inner circle: TE families and unknown repeats in the scaffolds (see the legend for details). Vertical axis of each inner circle: The mean distance of neighbouring genes/TEs. Short distances between the genes/TEs result in dots towards the centre of Circos plot whereas long distances result in dots towards the outer circle. Links: Syntenic regions shared.

**Figure 7.**
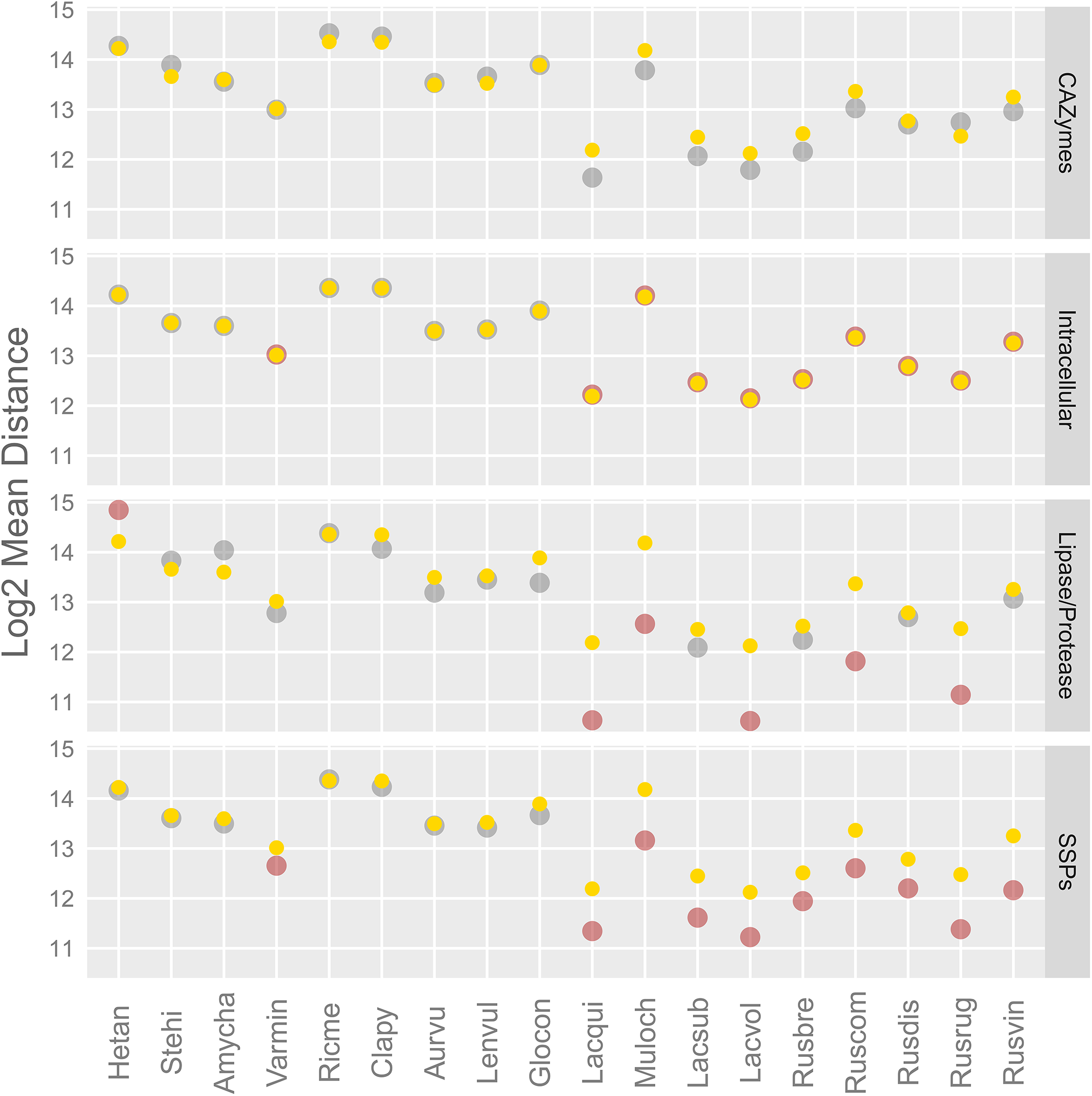
Mean distances between genes and TE clusters. Yellow: Mean distances averaged from the 5,000 reshuffled models. Red: Mean distances observed in the genomes with statistical significance (p < 0.01). Grey: Mean distances observed in the genomes (p > 0.01). Distances (base) are transformed in log2 (Table S14).

The concerted loss of CAZymes has been observed in conserved portions of Russulaceae genomes by comparing the largest syntenic region. This comparison reveals contrasting retention of the genes of interest (e.g., AA3_2, AA2, AA9 and GH74) between the saprotrophic *G. convolvens* and closely related ECM species that have lost these traits, such as *R. vinacea* or *R. brevipes* (Fig. 5). A conserved syntenic cluster of secreted CAZymes was detected as a core secretory capacity for Russulaceae, which includes glycosyltranferases and carbohydrate esterases. Secondary core capacity is widespread for a larger array of secreted proteins that includes glycoside hydrolases, polysaccharide lyases, expansin-like proteins, carbohydrate binding modules, and glycosyltransferases. Repeated content pockets are infrequent across the syntenic region and do not show a correlation with secreted genes, indicating that association between repeated elements and secreted genes are non-syntenic due to the activity of TEs.

### Impact of transposable elements on the genome landscape

The larger size of genomes of ECM Russulales species is mainly due to their higher content in repeated elements. TEs comprise a higher copy number and genome coverage (%) in ECM Russulaceae than in other Russulales (p<0.01, WSR test, 2-tailed) ranging from 29 to 67% of genome assemblies and 27,000 to 164,000 total copies (Fig. 1b; Fig. S2). *Gypsy, Copia* Long Terminal Repeat (LTR) retrotransposons, hAT families and other unclassified repeats are among the most abundant TEs in genomes from ECM Russulaceae (p<0.01, WSR test, 2-tailed). hAT repeats are involved in RNA processing and are unique to ECM Russulaceae (Hammani et al. 2012). Notably, *Penelope* non-LTR retrotransposons are only present in *L. quietus*.

ECM Russulaceae genomes show statistically significant associations between secreted proteins and TE rich areas with all secreted protein classes more closely clustered to TEs than non-ECM Russulales (Fig. 7). For non-secreted proteins, ECM Russulaceae genes are more distantly spaced from TEs than non-ECM Russulales. ECM Russulaceae show extreme clustering of TEs with secreted lipases/proteases and SSPs. Intracellular genes are isolated from TEs for all of ECM Russulaceae and *V. minispora*, which also shows the same trend of distance for SSPs and TEs. Distances between TEs and lipases/proteases show the opposite trend in *H. annosum*, a root pathogen, with genes being clustered with TEs.

## Discussion

### Evolutionary adaptation towards ectomycorrhizal lifestyle

Genomes of ECM Russulaceae fungi present many of the hallmarks of the transition to ECM symbiosis, including an expansion in genome size due to the accumulation of repeated elements and a contraction in gene families involved in the enzymatic breakdown of plant organic matter (Kohler et al. 2015; Martin et al. 2016; Hess et al. 2018; Miyauchi et al. 2020). The contraction of the total PCWDE gene repertoire coincides with the evolution of the ECM habit after the split with the closest extant saprotrophic species, *G. convolvens*. Across the pangenome, however, a heightened rate of gene evolution comprised of a high rate of gene loss and duplication was detected as preceding the evolution of the ECM habit. This can be seen in some PCWDE orthogroups but is also a general pattern across the pangenome. This trend was shown in the POD gene family contraction for the common ancestor of Amanitaceae, another group of mostly ECM species with closely related saprotrophs and a single switch to ECM (Kohler et al. 2015; Hess et al. 2018). Russulaceae provides evidence that this trend may be pervasive across the genome and an outcome of protracted genome remodeling for priming a switch to symbiosis. Protracted rates of gene gain, loss, and duplications across the ancestral pangenome of Russulaceae suggests that preadaptive priming to the ECM habit is likely tied to changes in the mode of nutrition (i.e., polysaccharide metabolism) instead of signaling pathways controlling biotic interactions (e.g., effector-like SSPs). These pre-existing traits may have emerged more frequently in facultative saprotrophs loosely interacting with tree roots with the ability to switch nutritional modes being latent across a wide diversity of fungi (Smith et al. 2017). The saprotrophic *G. convolvens* is known to widely colonize well-decayed logs using a suite of oxidases, which may necessitate the ability to circumvent plant root defenses within the wood for substrate occupation (Nakasone 1990). Some species of ECM Russulaceae form mushrooms on well-decayed logs which are thought to associate with roots within the wood and may utilize the POD ligninases retained from white rot ancestry to occupy this niche (Määkipä et al. 2017). Other elements of the secretome such as PCWDEs, FCWDEs, and proteases that saw modifications concurrent with the evolution of the ECM habit are potentially more essential to the ECM lifestyle for Russulaceae than host recognition pathways, effectors to circumvent host defense, and competitive interactions with other rhizospheric fungi.

Despite over 65 million years of divergence time between lineages of Russulaceae, species share a high degree of conserved gene similarity and synteny. Compared with the non-ECM Auriscalpiaceae, which shares an equivalent time of diversification, divergent lineages within Russulaceae maintain at least five times more syntenic links than members of Auriscalpiaceae. This high degree of gene order conservation may be essential for maintaining a conserved niche and lifestyle. While a large proportion of SSPs are species-specific, a conserved SSP was detected for Russulaceae coding for a phosphatidylglycerol/phosphatidylinositol transfer protein. A homolog of this protein was significantly accumulated in cork oak roots developing ectomycorrhizae with *Pisolithus tinctorius* (Sebastiana et al. 2017). This particular SSP may function as an effector to control the membranes of host plant cells for efficient nutrition exchange and hints at a latent mechanism for ECM symbiosis. This level of conservation is counter to the paradigm of effectors arising through convergent evolution and is quite uncommon (Kohler et al. 2015). No difference was detected in number of SSPs between saprotrophic and symbiotic members of Russulales.

The peculiar expansions of secreted protease and chitinase families may indicate a possible specialization of Russulaceae fungi to target non-plant derived organic sources of nitrogen such as fungal and bacterial necromass. Aspartyl proteases have been implicated in working in conjunction with hydroxyl radicals to access organic nitrogen from protein sources and have been detected as upregulated in the presence of soil organic matter for the ECM *Paxillus involutus* (Beeck et al. 2018; Shah et al. 2016). The most extreme gene expansion of 283 genes in a single orthologous cluster of subtilases with pro-kumamolisin activation domains was observed for *L. quietus*. Proteases encoded by these genes are involved in pathogenicity in animal and myco-parasitic fungi (Muszewska et al. 2011). *Russula gracillima* and *R. exalbicans* have been shown to parasitize *Lactarius* mycorrhizae (Beenken et al. 2004), but parasitism in Russulaceae is undoubtedly rare. An alternate explanation is that this pathway has been co-opted for plant host interaction. Additional sequencing of genomes in *Lactarius* will determine whether the whole genome expansion has taken place in the common ancestor of *Lactarius*.

We detected a reduction in the repertoire of gene clusters involved in secondary metabolism among the ECM Russulaceae, which has not been noted for an ECM lineage in any previous study. This may constitute another hallmark feature of the evolution of the ECM habit or may be specific to Russulaceae. This was particularly pronounced in NRPS-like (nonribosomal peptide-synthase) secondary metabolite gene clusters (SMCs), which have diverse functions but are most known for the production of mycotoxins and antibiotics (Bushley & Turgeon 2010). Russulaceae saw an expansion in siderophore SMCs containing conserved N-terminal iron uptake chelate (IucC) domains, which have been implicated in pathogenesis in *Rhizopus* and may be important for iron sequestration (Carroll & Moore 2018). Based on gene cluster similarity, Russulaceae possess a conserved set of terpene-related SMCs that are likely involved in the production of diverse lactarane sesquiterpenes that have frequently been characterized in Russulaceae (Clericuzio et al. 2012). An SMC of the “other” category conserved for Russulaceae was found to contain an aspartyl protease domain that may be part of the specialized function of Russulaceae. We hypothesize that a reduction in NRPS-like SMCs may correspond to a reduction in defensive compound diversity in ECM due to co-option of plant host defensive and a subsequent release on selective pressure.

### Functional specialization within Russulaceae

While the prevailing trend of a loss of PCWDEs in ECM lineages is evident in ECM Russulaceae, the counter trend of functional specialization within secreted CAZymes can be seen in different lineages of Russulaceae. Marked expansions in shared and unique homologous clusters of secreted enzymes as well as gene families indicates selection for specialized function in decomposition capability. ECM decomposition, such as “litter bleaching,” has been proposed as a significant contributor to carbon metabolism in forested ecosystems with phylogeny significantly predicting enzymatic activity (Talbot et al. 2008; 2015; Bödeker et al. 2014; Zak et al. 2019). The differential ability to scavenge nitrogen, phosphorus, and trace elements as key functions is mediated through these enzymes’ ability to break down soil organic matter, which can be detected by the plant host to mediate and select for its mycorrhizal community (Hortal et al. 2017). Traits that have been highlighted as potential drivers of diversification in ECM fungi have primarily looked at morphological traits of sexual reproduction and general ecological strategies, but adaptative functional specialization within ECM lineages is understudied and may be the key to understanding hyperdiversification and host dynamics within these groups (Looney et al. 2018; Sánchez-García et al. 2020). We hypothesize that niche partitioning is evident among ECM communities, with functional specialization followed by host switching driving diversification of ECM lineages. The extent to which differential ECM decomposition ability within ECM lineages is present and how these traits are partitioned within an ECM community should be further explored.

### Transposable elements driving gene innovations and ECM regulation

Expansions in TE content in the ancestor of Russulaceae may have facilitated adaptive shifts, such as loss of PCWDEs, expansion of protease/lipases, and diversification of SSPs involved in mycorrhiza development. This remodeling is inferred to have coincided with the K-Pg boundary extinction event, suggesting that the shift may have been driven by a drastic shift in plant community composition due to mass extinction(Nichols and Johnson 2008). ECM plant hosts at this time began a shift towards highly variable root evolution within plant families, and diversification of many angiosperm host species occurred later, during the early diversification of lineages within Russulaceae (Looney et al. 2016; Brundrett et al. 2018). An associated growth habit with plant roots of soil decomposers and lignicolous saprotrophs, the habit of *G. convolvens,* would have allowed for frequent interactions and coevolution eventually leading to the ECM lifestyle, and potentially, diversification.

Accumulation of repeated elements in the genomes of ECM Russulaceae and the close physical proximity of TE clusters and SSP genes suggest that TEs may promote gene innovation (e.g., promote duplication in SSPs, proteases) in ECM Russulaceae fungi. When TE insertions occur near host genes, expression is potentially altered due to the silencing of the TE through methylation mechanisms or TE activity on host cis-regulatory elements (Chuong et al. 2017). The proliferation of TEs in ECM Russulaceae might have led to the formation of TE hotspots that contribute to complex life traits, such as those involved in developmental signaling pathways. The patterns of localized TEs seem to be species-specific. Such localized TEs may have contributed to unique transcription regulations and gene expression. Our findings are consistent with the view that accumulation of TEs in particular genomic regions have affected certain genes that trigger morphological and physiological changes that are key to the ECM symbiosis (Chuong et al. 2017; Sultana et al. 2017).

### Conclusion

In some lineages, such as Russulaceae and Amanitaceae to some extent, genetic traits typifying the evolution of ECM fungi (e.g., loss of PCWDE orthologs, expansion of TE content; reduction of SMCs) are already observed in the genomes of closely related saprotrophic species, and this pre-existing trait may explain the pervasive, recurrent evolution of ECM associations. While the evolution of the ECM habit releases selection on genes required to access plant carbon in the soil, these genes can be coopted for functional specialization in the fungus’s ability to access nutrients, colonize the apoplastic space of the host roots, and/or gain a competitive advantage during community assembly. This specialization may be tightly linked to co-evolutionary host-interactions, mediated by a heightened adaptability of ECM fungi through a heightened rate of gene expansion and turnover through TE association. Whole-genome sampling within Russulaceae targeted single representatives of a highly diverse group, so additional sequencing of targeted groups will help to test hypotheses of functional specialization and its relationship to diversification.

## Materials and Methods

### Taxon sampling and nucleic acid extraction

Newly sequenced genomes and transcriptomes were derived from phylogenetically distinct lineages within the family Russulaceae according to Weiß et al. (2016). Representative species were sampled as mushroom sporocarps from forested habitat in the Great Smoky Mountains National Park and surrounding areas. To retrieve high molecular weight DNA and undegraded RNA, the inner flesh of the sporocarps was extracted in the field using a sterilized scalpel and placed in a 50 mg Falcon tube. Material was then flash-frozen in the field in liquid nitrogen. Tissue samples were also attempted on Melin-Norkrans Modified media with collections for experimental applications with a low success rate. A member of the closest related extant outgroup, *G. convolvens*, was also sampled for comparative analyses between different trophic modes. Vouchered specimens are accessioned in the herbarium of the University of Tennessee.

Extraction of high molecular weight DNA was performed using a cetyltrimethylammonium bromide (CTAB) based protocol. Frozen sporocarp material was first ground into a fine powder in liquid nitrogen using a mortar and pestle. The powder was added to 1.5 mL Eppendorf tubes weighed at 90 mg increments. A pre-warmed (~55° C) lysis buffer was added to each sample at a volume of 700 μL. The lysis buffer consisted of a mixture of 260 μL of buffer A (0.35 M sorbitol, 0.1 M Tris HCl pH 9, and 5 mM EDTA pH 8), 260 μL of buffer B (0.2 M Tris HCl pH 9, 50 mM EDTA pH 8, 2M NaCl, and 2% CTAB), 104 μL buffer C (5% Sarkosyl [N-lauroylsarcosine sodium salt], and 70 μL of a 0.1% solution of Polyvinylpyrrolidone (PVP). Samples were then centrifuged at 14 000 rpm for 3 minutes to compact rehydrated biomass. A micropestle was used for additional grinding and this process was repeated at least one more time. Protein digestion was performed by adding 5 μL of Proteinase K (10mg/mL), vortexing, and incubation of samples for 30 min. at 65° C. For sodium dodecyl sulfate precipitation, 230 μL of 5 M KAc was added to samples, inverted to mix, and incubating for at least 30 minutes in ice or for 16 hours in a 4° C refrigerator. Following incubation, samples were centrifuged at 14 000 rpm for 10 minutes and 1 mL of supernatant was transferred to 2 mL Eppendorf tubes. An equal volume of Chloroform:Isoamylalcohol (24:1) was added and the tubes were centrifuged for 10 minutes at 14 000 rpm. A conservative amount of supernatant (~850 μL) was drawn avoiding the top and bottom layers and added to additional 2 mL Eppendorf tubes. Again, an equal amount of Chloroform:Isoamylalcohol (24:1) was added and centrifuged for 10 minutes. A final volume of 675 μL was added to new 1.5 mL Eppendorf tubes and treated with an RNase digestion with 10 μL of RNAseA (100mg/mL) and incubated at 37° C for 10 minutes. DNA precipitation was done by adding 67.5 μL of 3 M NaAc pH 8 and 675 μL of absolute isopropanol and incubated for 5 minutes at room temperature. The samples were then centrifuged for 10 minutes at 4° C and the surnageant was eliminated by gently pouring it off. Ethanol washing was done with 200 μL 70% ethanol followed by centrifugation at 14 000 rpm at 4° C. Ethanol was then carefully drawn out using a double pipet tip method making sure not to disturb the pellet. Samples were then dried for 5 minutes in a vacuum pump to completely dry the pellet. The pellets were then resuspended in 10 μL of TE buffer and stored at 4° C for quality assessment.

Extraction of RNA was performed using a Sigma™ Plant Total RNA Kit. Surfaces were first sterilized with 70% ETOH and D/RNAse Free™ decontaminant to prevent enzyme contamination. Frozen sporocarp material was again ground into a fine powder in liquid nitrogen using a decontaminated mortar and pestle. The powder was added to enzyme-free 1.5 mL Eppendorf tubes weighed at 100 mg increments. The provided lysis buffer was added to each sample at a volume of 500 μL. Samples were then centrifuged at 14 000 rpm for 3 minutes to compact rehydrated biomass. A micropestle was used for additional grinding and this process was repeated at least one more time. Once samples were sufficiently ground, 5 μL of 2-mercaptoethanol was added to each sample and incubated at 55° C. The rest of the protocol followed the provided protocol of the kit, using Protocol A for the binding step and following the optional On-Column DNase Digestion procedure. Once product was eluted, 1 μL of Roche Protector RNase Inhibitor was added to stabilize the product. An aliquot of 9 μL was stored at 4° C° for quality control and the remaining sample was stored at −80° C.

Quality assessment followed the recommendations of JGI for DNA and RNA. First, nucleic acids were visualized using gel electrophoresis on a 1% agarose gel with Roche DNA Molecular Weight Marker II as ladder. Bands were evaluated based on brightness, amount of smearing, and presence or absence of contamination (i.e. RNA or DNA). Concentrated and undegraded DNA samples were pooled after centrifugation at low speed for one minute to homogenize and without pumping the pipet. Assessment of concentration and total amount for genomic DNA was assessed using the Qubit^®^ DNA BR Assay Kit on a Qubit^®^ 2.0 fluorometer. Assessment for RNA concentration and quality was done using an Experion™ RNA Analysis kit analyzed using the Experion™ Automated Electrophoresis System. RNA with clear bands that achieved an RQI score of at least 6.5 was deemed adequate for JGI submission.

### Genome sequencing, assembly, and annotation

All sequencing, assembly, and annotation was performed at JGI. Genome sequencing was done with Pacific Biosciences (PacBio) technology, except for 4 genomes sequenced with Illumina technology. All PacBio-sequenced genomes were assembled using Falcon (Chin et al., 2016). Of the Illumina-sequenced genomes, 3 were assembled serially using Velvet (Zerbino and Birney 2008) followed by ALLPATHS-LG (Gnerre et al., 2011), and 1 was assembled with ALLPATHS-LG alone. Mitochondrial genomes were assembled separately. All transcriptome sequencing was done with Illumina only, and subsequently assembled into putative transcripts using Trinity (Grabherr et al., 2011) or Rnnotator (Martin et al 2010). Each genome was annotated using the JGI Annotation Pipeline (Grigoriev et al., 2014; Kuo et al., 2014) aided by the transcriptome. Detailed methods are described in Appendix 1 of the Supplementary Materials.

### Comparative genomic feature analyses

Statistics of JGI genome assemblies (i.e., N50, number of genes and scaffolds, genome size) were obtained from JGI MycoCosm (https://mycocosm.jgi.doe.gov). Genome completeness with single copy orthologues was calculated using BUSCO v3.0.2 with default parameters (Simão et al. 2015). The TE coverage in genomes was calculated using a custom pipeline Transposon Identification Nominative Genome Overview (TINGO; Morin et al. 2019). The information above was combined and visualized. Secretomes were predicted as described previously (Pellegrin et al. 2015). We calculated, visualized, and compared the count and ratio of total (present in the genomes) and predicted secreted CAZymes, lipases, proteases, and small secreted proteins (< 300 amino acid) as a subcategory. We calculated the total count of the followings using both all and predicted secreted plant cell wall degrading enzymes (PCWDEs) and microbe cell wall degrading enzymes (MCWDEs). Global trends of ecological groups were evaluated using Non-metric Multi-Dimensional Scaling (NMDS) with the count of total and predicted secreted CAZymes. The dissimilarities among the ecological groups were calculated and the relationship was converted into distances in the two-dimensional space with the function mataMDS in R package Vegan (Oksanen et al. 2016). We grouped fungi into broad ecological categories and assessed secretome differences between the ecological groups by performing non-parametric multiple comparisons with the function gao_cs in R package nparcomp (Konietschke 2009). We examined the total and predicted secreted counts of CAZymes/ lipases/ proteases/ SSPs. Statistically significant ecological groups (p < 0.05) were determined. Output files generated above were combined and visualized with custom R scripts, Proteomic Information Navigated Genomic Outlook (PRINGO; Miyauchi et al. 2020).

### Phylogenomic inference and molecular clock analyses

We constructed a phylogeny based on orthologous genes among the selected fungi using FastOrtho with the parameters set to 50% identity, 50% coverage, inflation 3.0 (Wattam et al. 2014). Protein sequences used for the process were genome-wide protein assemblies from JGI fungal portal MycoCosm. We identified clusters with single copy genes, aligned each cluster with MAFFT 7.221 (Katoh et al. 2013), eliminated ambiguous regions (containing gaps and poorly aligned), and concatenated single-gene alignments with Gblocks 0.91b (Castresana 2000). We constructed a phylogenetic tree with RAxML 7.7.2 (Stamatakis 2014) using the standard algorithm, the PROTGAMMAWAG model of sequence evolution and 1000 bootstrap replicates.

A set of 38 genomes across the Agaricomycotina were selected for calibrating a molecular clock and dating of the Russulales lineage. Gene selection for molecular clock analysis of Russulales was done based on phylogenetic informativeness performed in PhyDesign (López-Giráldez & Townsend 2011). Molecular clock analysis was performed in BEAST v1.8.4 (Drummond et al. 2012) using the 20 most phylogenetically informative loci due to computational constraints of the program for dealing with large datasets. Three calibrations based on fossils were used: *Archaeomarasmius leggetti*, an agaric fossilized in 90 Ma Dominican amber as the minimum age of Agaricales (Hibbett et al. 1997); *Quatsinoporites cranhamii*, a poroid shelf fungus estimated at 113 Ma as the minimum age of the Hymenochaetales (Smith et al. 2004); and *Geastroidea lobata,* a gastroid fruiting body with a double-layered peridium from the Cretaceous (72–66 Ma)(Krassilov and Makulbekov 2003). The analysis used an uncorrelated lognormal relaxed clock model prior with a Constant Coalescent tree prior. MCMC was run independently three times for fifty million generations, logging every 1000 generations. The runs were checked for convergence and mixing using Tracer v1.6 (Rambaut et al. 2013). An ultrametric maximum-clade-credibility (MCC) tree was summarized in TreeAnnotator 1.8.4 with a burn-in of 25% of trees.

### Secondary metabolite analysis

Secondary metabolite clusters (SMCs) were predicted using antiSMASH 5.0 using a relaxed strictness through the online dedicated server (Blin et al. 2019). Filtered gene models were used as feature annotations. Resulting .gbz files were analyzed through the BiGSCAPE pipeline using default parameters and the Pfam-A v30.0 database (Navarro-Muñox et al. 2020).

### Genome rearrangement analysis

Syntenic blocks were identified from pair-wise comparisons of genomes with R package DECIPHER (Wright 2015). Macrosynteny was determined using “FindSynteny” function with default parameters with the argument for masking repeat sequence turned off whereas mesosynteny was identified using with the modified parameters (i.e. maxSep = 1000, maxGap = 1000) suitable for highly similar sequences at the gene level. We made the genomic coordinates of genes from JGI genome-wide gene catalogue (Grigoriev et al. 2014). JGI functional gene annotations from the InterPro database were used for the description of intracellular and extracellular proteins (https://mycocosm.jgi.doe.gov). Then, it was combined with the secretome and repeats data described above. The integrated results were used for the circular representation of the genome assemblies with the combined genomic information using R package circlize (Gu et al. 2014). Also, we measured the mean TE-gene distances with statistical support by comparing the locations of observed genes and TEs and 5,000 null hypothesis genome models made by randomly reshuffling the locations of genes. The probability (p value) of mean TE-gene distances was calculated with R package, regioneR (Gel et al. 2016). The process above was conducted with a set of custom R scripts, Synteny Governance Overview (SynGO; Hage et al. 2021). Scaffolds containing major syntenic regions among the species were visualized along with the identified predicted secretome and TEs using R package karyoploteR (Gel & Serra 2017). Data integration was performed with a set of custom R scripts, Visually Integrated Numerous Genres of Omics (VINGO; available upon request).

### Gene evolution analysis

Evolutionary gains or losses of orthologous gene groups were estimated on the basis of the constructed phylogeny using Software for Computational Analysis of gene Family Evolution (CAFE; De Bie et al. 2006). The software uses a random birth and death process to model gene gain and loss across a user specified tree structure. The distribution of family sizes generated under the random model provides a basis for assessing the significance of the observed family size differences among taxa. We selected gene families with p value < 0.001.

### Evolutionary rate analysis

Orthologous gene clusters were imported into COUNT (Csurös et al. 2018) for genome and secretome clustering analyses to assess gene evolution rates and reconstruct gene family history. Gene clusters containing fewer than 3 species were filtered out of the rate optimization. Rate optimization used the Gain-loss-duplication model with a Poisson distribution at the root and lineage-specific variation estimated. The analysis was run for 100 rounds with a convergence threshold on the likelihood of 0.1. Gene ancestral reconstruction was inferred using Dollo parsimony and posterior probabilities using a birth-and-death model.

## Data availability

The genome assemblies used for this study are available on the JGI fungal genome portal, MycoCosm (https://mycocosm.jgi.doe.gov). See the details of JGI genomes used for the study including DDBJ/ENA/GenBank accession numbers (Table S15). The latest CAZyme annotations are available upon request from CAZy team, Aix-Marseille University, France.

## Acknowledgments

This research was supported by the Genomic Science Program, U.S. Department of Energy, Office of Science, Biological and Environmental Research as part of the Plant Microbe Interfaces Scientific Focus Area, at the Oak Ridge National Laboratory. Oak Ridge National Laboratory is managed by UT-Battelle, LLC, for the U.S. Department of Energy under contract DE-AC05-00OR22725. The work conducted by the U.S. Department of Energy Joint Genome Institute, a DOE Office of Science User Facility, is supported by the Office of Science of the U.S. Department of Energy under Contract No. DE-AC02-05CH11231. This was done within the framework of the Community Sequencing Program #1974 “*1KFG: Deep Sequencing of Ecologically-relevant Dikarya”* and the Community Sequencing Program #305 “*Mycorrhizal Genomics Initiative*”. Research in the Martin laboratory is also funded by the Laboratory of Excellence Advanced Research on the Biology of Tree and Forest Ecosystems (ARBRE; grant ANR-11-LABX-0002-01), the Region Lorraine Research Council and the European Commission (European Regional Development Fund). BL would like to thank the Laboratory of Excellence ARBRE for a visiting researcher grant. Thanks to Daniel Lindner at the USFS Northern Research Station and Otto Miettinen at the University of Helsinki for taxonomic expertise and providing sample support. We would like to thank Prof. Joseph Spatafora and Rytas Vilgalys for their support.

## Author contributions

BPL, JL, PBM and FM conceived the project. FM coordinates the Mycorrhizal Genomics Initiative. IVG and KWB coordinated the genome sequencing, assembly and annotation at JGI. SM, BPL, AK and EM performed comparative genome analyses. ED and HB provided updated cazy profiles. BPL, DH, LN, and PEC extracted DNA and RNA. EL, AL, KL, AK, MN, AT carried out DNA sequencing, RNA sequencing, genome assembly or gene prediction at JGI. BPL, SM, FM wrote and edited the manuscript. All co-authors read and edited the manuscript.

## Materials & Correspondence

Please contact the corresponding authors Brian Looney or Francis Martin.

