## Supplementary material for "Evolutionary priming and transition to the ectomycorrhizal habit in an iconic lineage of mushroom-forming fungi: is preadaptation a requirement?": Figures S1-17

**Supplementary Figures**

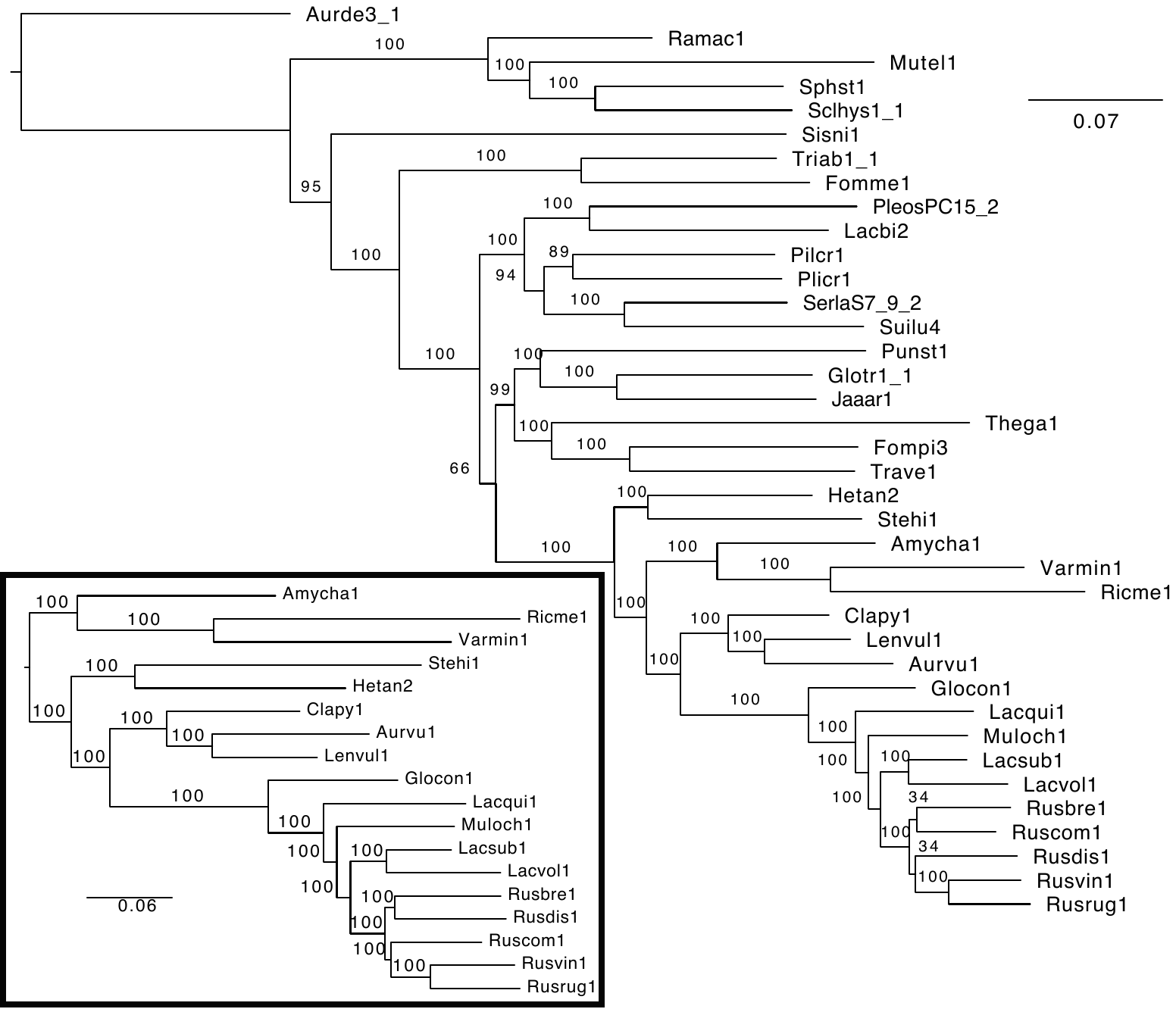

***Figure S1. Phylogenomic reconstruction of Russulales and Russulales under Maximum Likelihood criterion****.* ***a.*** *Reconstruction of Agaricomycetes.* ***b.*** *Reconstruction of Russulales.*

An initial phylogenomic analysis of Agaricomycetes, based on 725 conserved single copy genes (532,877 amino acid positions) recovers high support for relationships between genera of Russulaceae, with *Lactarius* as sister clade to the rest of ECM Russulaceae, and *Lactifluus* as the sister clade to *Russula*. Relationships between major clades of *Russula*, however, are unresolved in this analysis. A focused phylogenomic analysis of Russulales taxa based on 2023 conserved single copy genes (1,300,326 amino acid positions) resolves the Russulales phylogeny with full bootstrap support at all nodes. Within *Russula*, *R. brevipes* and *R. dissimulans* are resolved as sister to the rest of the genus. The common ancestor of the ECM habit in Russulaceae is inferred to have arisen around the Cretaceous-Palaeogene (K-Pg) extinction event (73.6-60.1 MY), a period of rapid ecological and anatomical inovation (Alfaro et al., 2018).

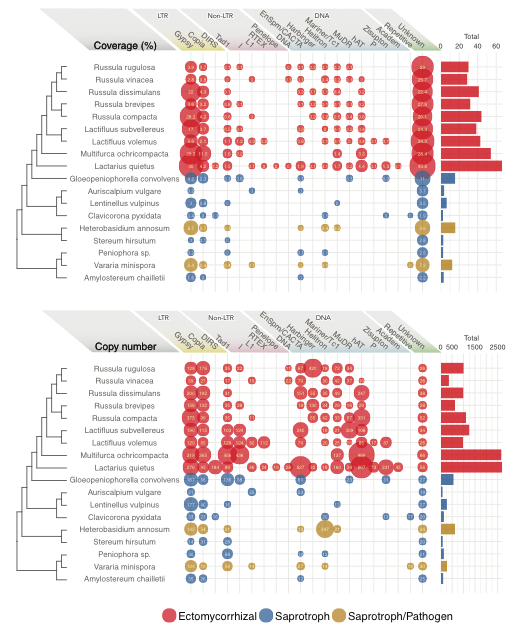

***Figure S2****.* ***Coverage of transposable elements identified****.* ***LTR****: Long terminal repeat retrotransposons.* ***Non-LTR****: Non- long terminal repeat retrotransposons****. DNA****: DNA transposons.* ***Repetitive****: Simple repeats.* ***Unknown****: Unclassified repeated sequences. The bubble size is proportional to the coverage of each of transposable elements (showing inside the bubbles). The right bars show the total coverage/ copy number per genome. See supplementary information for details (Table S# - 18Russulales_Step5_Taxo_Coverage_States.csv).*

Repeated elements comprise a higher copy number and genome coverage (%) in ECM Russulaceae than in other Russulales (*p*<0.01, WSR test, 2-tailed) ranging from 29 to 67% of genome assemblies and 27,000 to 164,000 total copies (Fig. S2). *Gypsy*, *Copia* Long Terminal Repeat (LTR) retrotransposons, *hAT* families and unknown repeats are highly covered in genomes from ECM Russulaceae (*p*<0.01, WSR test, 2-tailed), with *hAT* repeats involved in RNA processing^13^ and being unique to ECM Russulaceae.*Tad1* and *Mariner/Tc1* DNA transposons are found for all ECM Russulaceae genomes, but they can also be found for other Russulales genomes. *Gloeopeniophorella convolvens* possesses a larger proportion of overall TE content compared to other Russulales saprotrophs along with the two, putative dual saprotroph/pathogen species *H. irregulare* and *Vararia minispora*. This pattern also holds for Unknown TE classes, *Gypsy*, and *Copia* LTR. Non-LTR retrotransposons, *Penelope*, are present only in *L. quietus*.

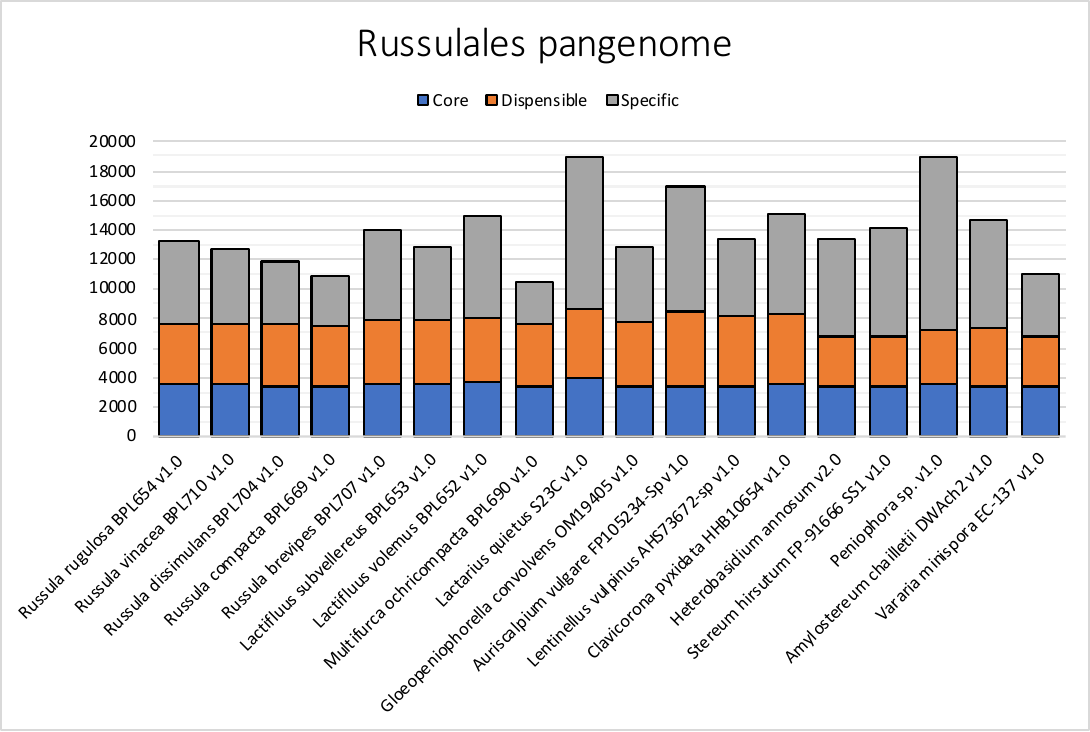

***Figure S3.*** *Pangenome of 18 Russulales. The number of genes is shown on Y-axis. Core: Common orthologous genes shared all species. Dispensable: Orthologous genes shared by* at least two species. *Specific: Species-specific genes.*

The pangenome of Russulales comprises over 250,000 gene models for all eighteen species, ranging from 10,514 genes for *M. ochricompacta* to 18,952 genes for *Peniophora* (Fig. S3). The core genes, including orthologous clusters missing from one or fewer species, make up one quarter of all genes ranging around 3500 genes for most species and up to 4023 genes for *L. quietus*. Dispensable genes (i.e. at least two species share orthologues) make up 30% of all genes with only slight differences in numbers between species. The species-specific gene content varies considerably between species but not trophic categories, with *Peniophora* sp. and *L. quietus* having the highest number of orphan genes (11721 and 10313 respectively) and *M. ochricompacta* with only 2832 orphan genes. Secondary alleles from dikaryotic genomes for ECM Russulaceae are detected to comprise 14-39% of all protein models (Table S# - Genome assemblies and annotation statistics).

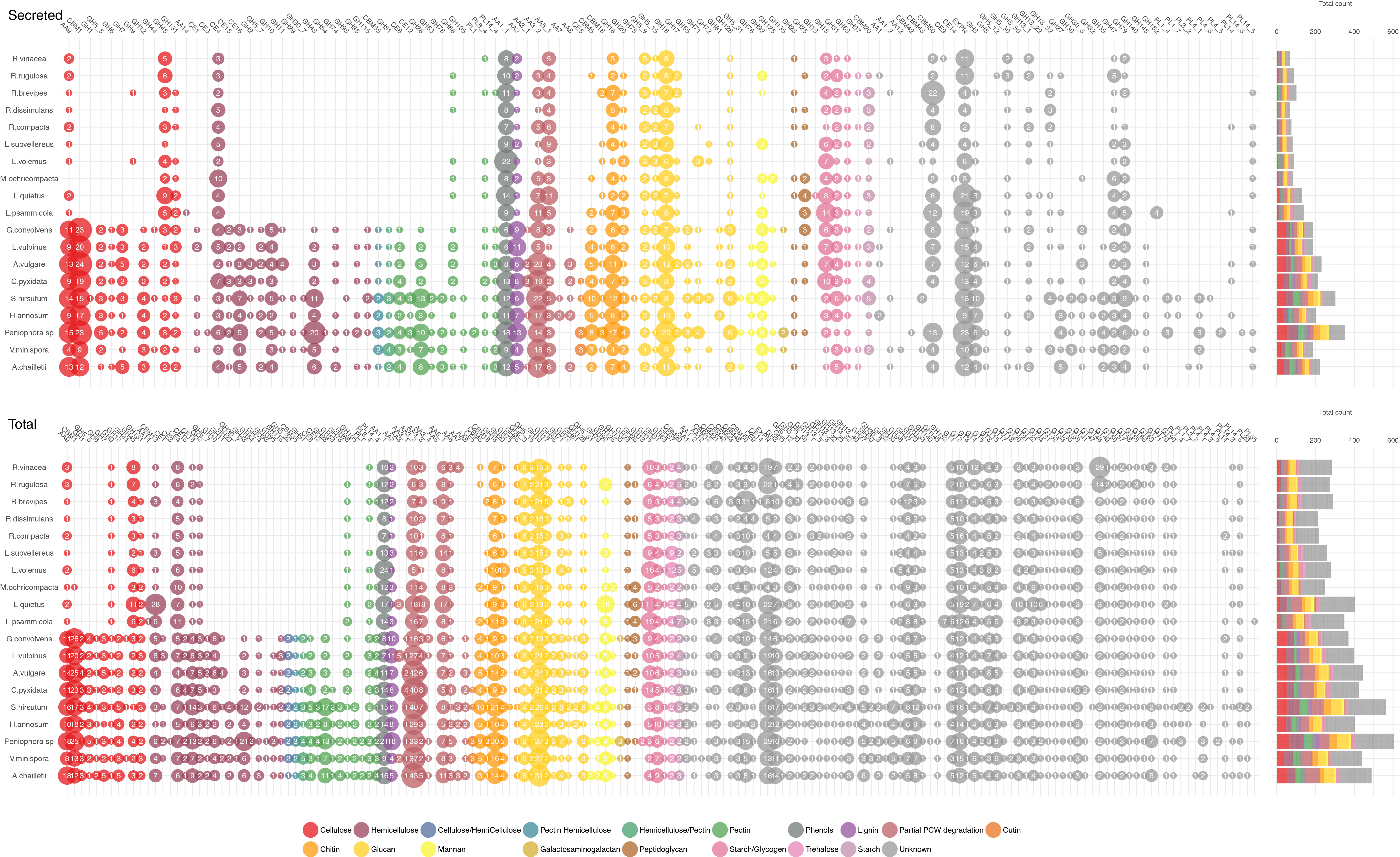

***Figure S4****.* ***Presence and absence of CAZyme domains for 19 fungi****. The size of bubbles corresponds to the count of CAZyme domains.* ***Secreted*** *(top): Theoretical secretomes.* ***Total*** *(bottom): The total of CAZyme domains found in the genomes. Substrate specificity is shown in colours. The fungi are in the evolutionary order. Total count per genome is shown on the right.*

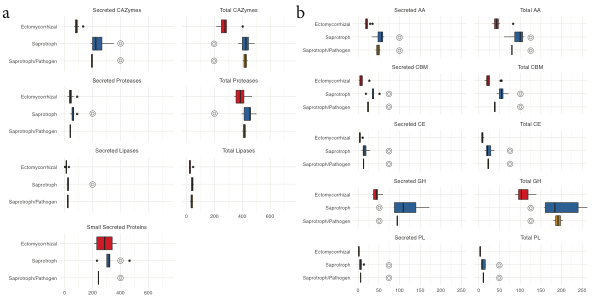

***Figure S5****.* ***Distributions of counts of theoretical secretome. a.*** *Lifestyle group-wise counts of theoretical secretome and total proteins present in genomes****. b.*** *Lifestyle group-wise count of theoretically secreted and total CAZyme families present in genomes. Double circles indicate statistically significant difference for the ectomycorrhizal group compared to the other ecological groups (p < 0.05). See supplementary table for details (Table S# - CAZy Lipase Protease SSP ANOVA; Table S# - CAZyme family ANOVA).*

We investigated lifestyle-wise distributions of counts for theoretically secreted and total proteins, and statistically evaluated if ECM fungi are different from any other groups. First, we assessed three protein categories (CAZymes, lipases, proteases) with a subcategory of small secreted proteins (< 300 aa; Fig S5a). Secondly, we sorted CAZymes based on the families (Fig S5b).

ECM fungi showed a significantly small number of secreted/total CAZymes, and secreted/total proteases compared to the saprotrophs (Fig S5a). When we examined CAZymes in details, the number of CAZyme families (AAs, CBMs, CEs, GHs, PLs) in ECM fungi was much smaller than those of saprotrophs and saprotropic pathogens (Fig S5b). It suggests ECM fungi have less secreted CAZymes, lipases, and protease compared to the saprotrophic groups. But, ECM fungi contain the higher proportion of genes for small secreted proteins (Fig 2). The distinctive pattern of ECM fungi could be explained by the plant dependent lifestyle.

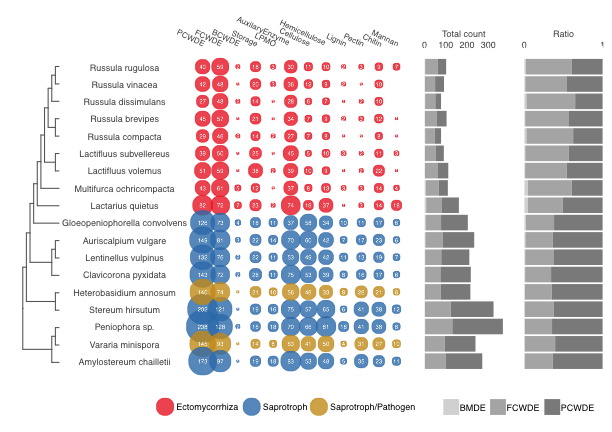

***Figure S6. Presence and abundance of CAZyme groups coded in genomes.*** *All the genes present in the genomes are counted (i.e. theoretically secreted and non-secreted). The fungi were labelled according to their ecology.* ***Bubble plot****: The number of including plant cell wall degrading enzymes (PCWDE) and microbial cell wall degrading enzymes (MCWDE), bacterial membrane (i.e. peptidoglycan) degrading enzyme (BMDE), lytic polysaccharide monooxygenase (LPMO), enzymes for starch and glycogen (storage); AA family CAZymes (Auxiliary Enzymes); substrate-specific enzymes for cellulose, hemicellulose, lignin, and pectin (plant cell walls); chitin, glucan, mannan (fungal cell walls).* ***Bar plots****: The total count of genes including PCWDE, FCWDE, and BMDE (left); and the proportion of PCWDE, MCWDE, and BMDE (right). See supplementary tables for counts and CAZyme list (Table S# - 18Russulales CEDE Step1 All Count&Stats.csv; 18Russulales_CWDE_Step1_Substrate_group_CAZyme_list.csv).*

We investigated secretome trends of 18 Russulales fungi. We compared the count of theoretically secreted and total coded CAZyme domains, lipases, proteases, others, small secreted proteins (Fig 2; Fig S6). CAZyme domains are separated into various categories. The ECM fungi contained the smaller number of secreted CAZymes and PCWDEs than the saprotrophic group (Fig 2). The ECM group lacks CAZyme families for degrading plant cell wall components (Fig S6).

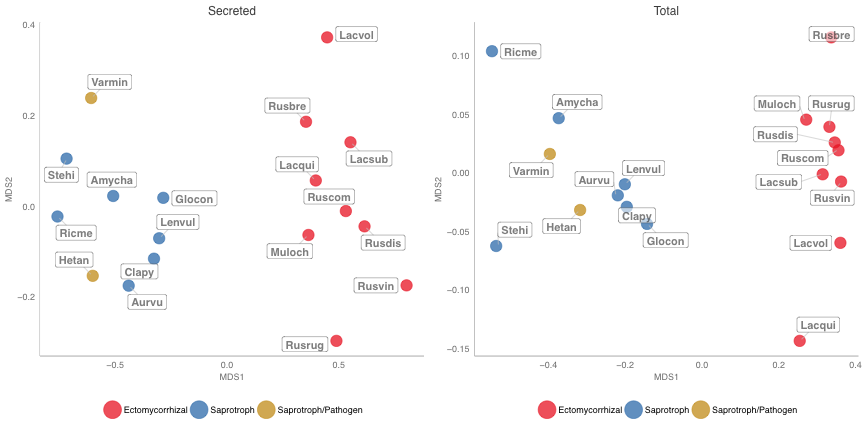

***Figure S7. Clusters of 18 fungi based on total and secreted CAZymes.*** *The fungi were grouped according to the number of CAZyme domains present in the theoretical secretome and the genomes. The colours represent their ecology. The labels show short JGI fungal IDs. Rusvin (R. vinacea), Rusrug (R. rugulosa), Rusbre (R. brevipes), Rusdis (R. dissimulans), Ruscom (R. compacta), Lacsub (L. subvellereus), Lacvol (L. volemus), Muloch (M. ochricompacta), Lacqui (L. quietus), Glocon (G. convolvens), Lenvul (L. vulpinus), Aurvu (A. vulgare), Clapy (C. pyxidata), Stehi (S. hirsutum), Hetan (H. annosum), Ricme (Peniophora sp.), Varmin (V. minispora), Amycha (A. chailletii). See supplementary tables for CAZyme domain counts (CAZyme tables in Count for total+secreted proteins).*

The count of theoretically secreted and total CAZymes coded in the genomes were transformed into two dimensional distances using Non-metric Multi Dimensional Scaling (NMDS; Fig S7). We found a global pattern in the grouped fungi. The first dimension separated fungi into two ecological groups, ECM fungi and saprotrphs including saprotrophic pathogens. Then, the second dimension spread out the species within the groups.

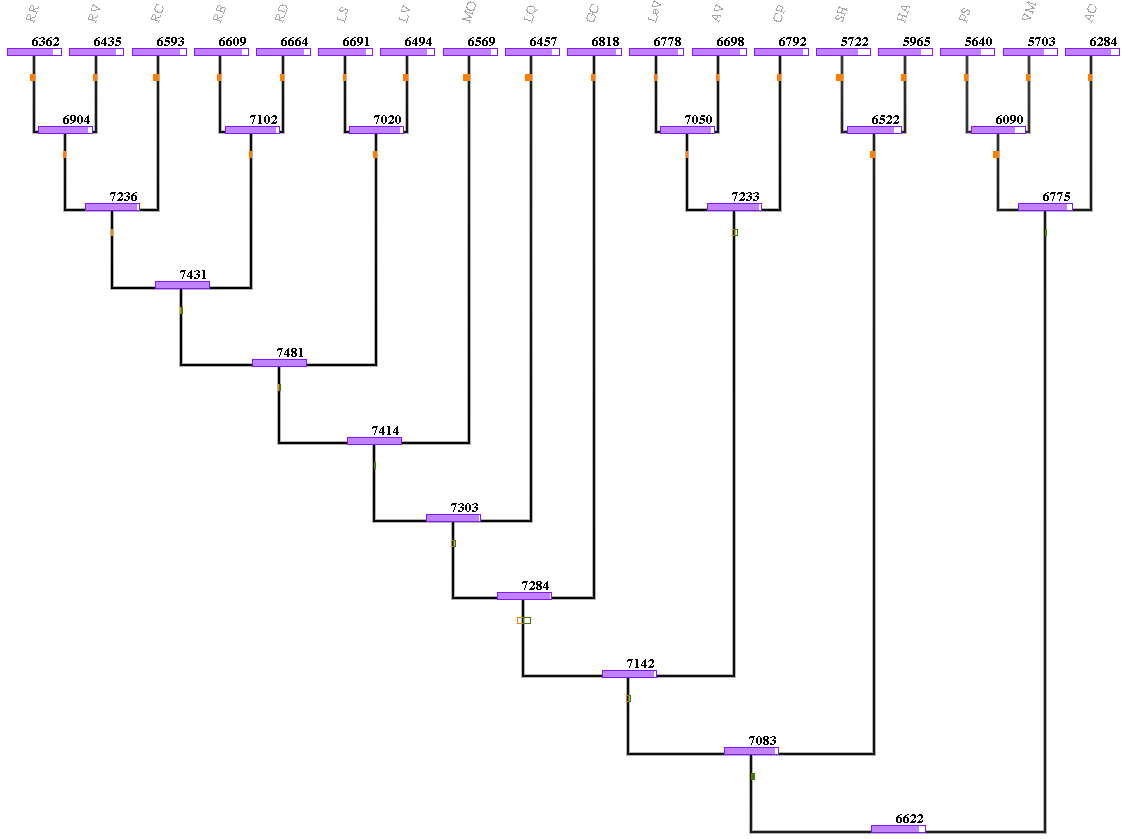

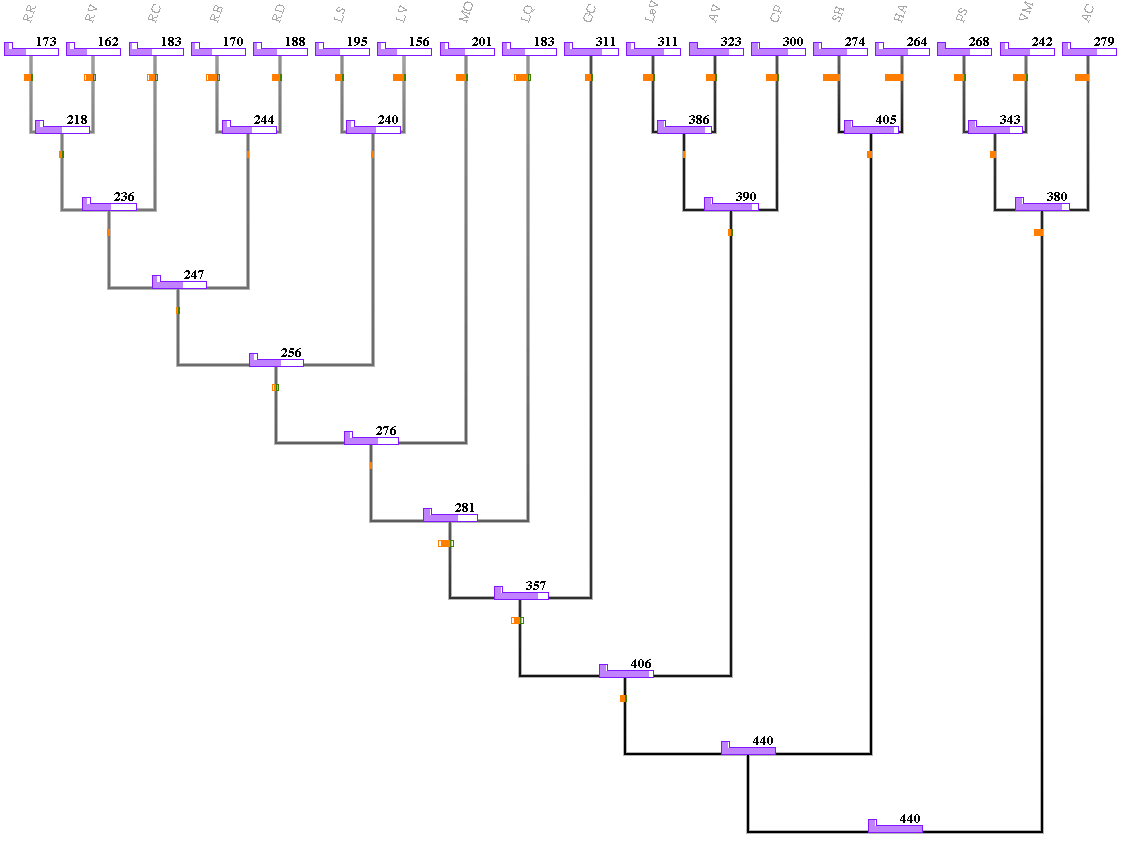

***Figure S8****.* ***Evolutionary rate COUNT analysis of Russulales. left.*** *Dollo parsimony reconstruction of ancestral pangenome content. Net gain (green) and loss (orange) are given as bars under nodes with horizontal shading indicating proportional likelihood.* ***right.*** *Dollo parsimony reconstruction of ancestral secretome content.*

Dollo parsimony reconstruction results of recapitulate results based on rates of gene gains, losses, and duplications within the Russulales. Net gain and loss occurred at a high rate in the ancestor of Russulaceae across the pangenome with species-specific net losses pronounced throughout the order. Reconstruction of the ancestral secretome reveals these same pronounced net gains and losses spread between the ancestor of Russulaceae and the ancestor of ECM Russulaceae.

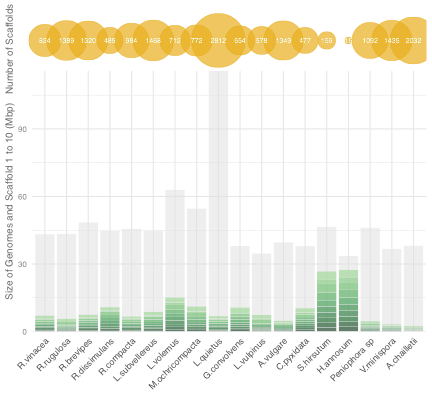

***Figure S9****.* ***Genome size and*** *s****caffold number****. The moon bamboo plot summarises the sequence quality of the 18 genomes.* ***Grey shadow****: the total size of scaffolds* ***Green sections****: the size of scaffold 1 to 10.* ***Yellow circles****: Number of scaffolds.*

We visualised the number and size of scaffolds compared with the genome size in order to examine the completeness of genome sequences (Fig S9). Scaffolds 1 to 10 of *S. hirsutum* and *H. annosum* constitute most of their genomes, indicating their chromosomes are almost complete. The rest of 16 genomes are rather fragmented as the first 1 to 10 scaffolds do not cover 50 % of the entire genomes.

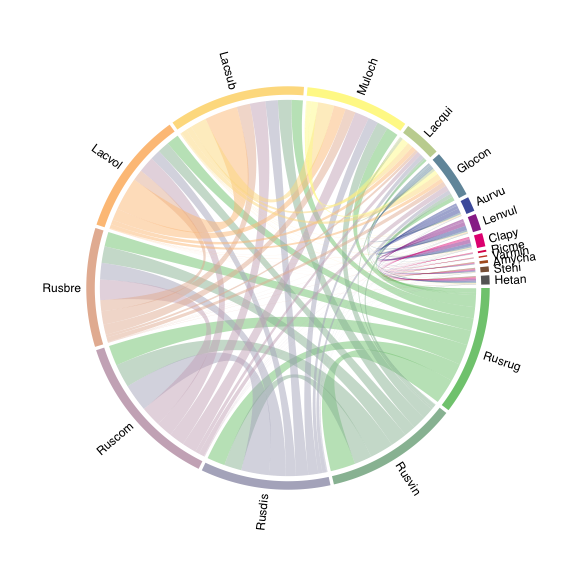

***Figure S10.*** *Overall synteny comparison of 18 Russulales. Number of shared synteny blocks among scaffold 1 to. Length of the bands with the fungal IDs represents the proportion of syntenic blocks. Links connecting the bands show syntenic blocks among the fungi. Rusvin (R. vinacea), Rusrug (R. rugulosa), Rusbre (R. brevipes), Rusdis (R. dissimulans), Ruscom (R. compacta), Lacsub (L. subvellereus), Lacvol (L. volemus), Muloch (M. ochricompacta), Lacqui (L. quietus), Glocon (G. convolvens), Lenvul (L. vulpinus), Aurvu (A. vulgare), Clapy (C. pyxidata), Stehi (S. hirsutum), Hetan (H. annosum), Ricme (Peniophora sp.), Varmin (V. minispora), Amycha (A. chailletii).*

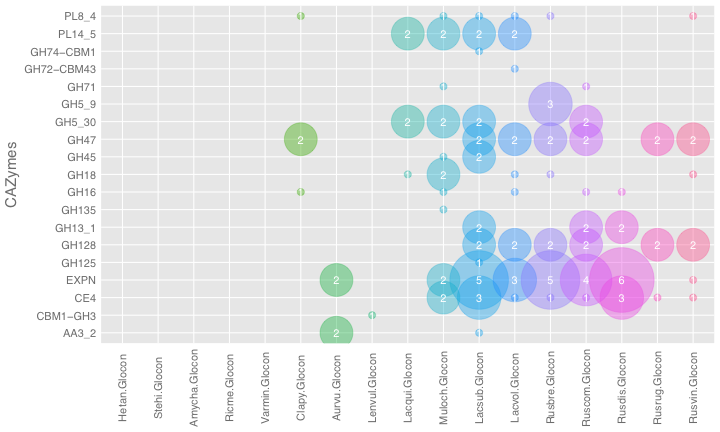

***Figure S11****.* ***Number of CAZyme-coding genes in synteny****. The count of CAZymes in the shared syntenic regions between G. convolvens (Glocon) and the rest of 17 Russulales fungi. Rusvin (R. vinacea), Rusrug (R. rugulosa), Rusbre (R. brevipes), Rusdis (R. dissimulans), Ruscom (R. compacta), Lacsub (L. subvellereus), Lacvol (L. volemus), Muloch (M. ochricompacta), Lacqui (L. quietus), Glocon (G. convolvens), Lenvul (L. vulpinus), Aurvu (A. vulgare), Clapy (C. pyxidata), Stehi (S. hirsutum), Hetan (H. annosum), Ricme (Peniophora sp.), Varmin (V. minispora), Amycha (A. chailletii).*

Secreted proteins encoded in shared syntenic regions between ECM Russulaceae species and *G. convolvens* include many expansins (EXPN), glycoside hydrolases (GH47, GH129), and a carbohydrate esterase (CE4) (Fig. S11). In most of Russulales, the highly conserved syntenic regions also include genes encoding ATPases, protein kinases, WD40 repeat-containing proteins, zinc finger transcriptional factors, suggesting these genes are essential for maintenance of cellular function. Russulaceae*-*specific genes are coding for Zn(2)-C6 fungal-type NDA binding domains, DEAD/DEAH box helicases, and RNA recognition motifs (Fig S12).

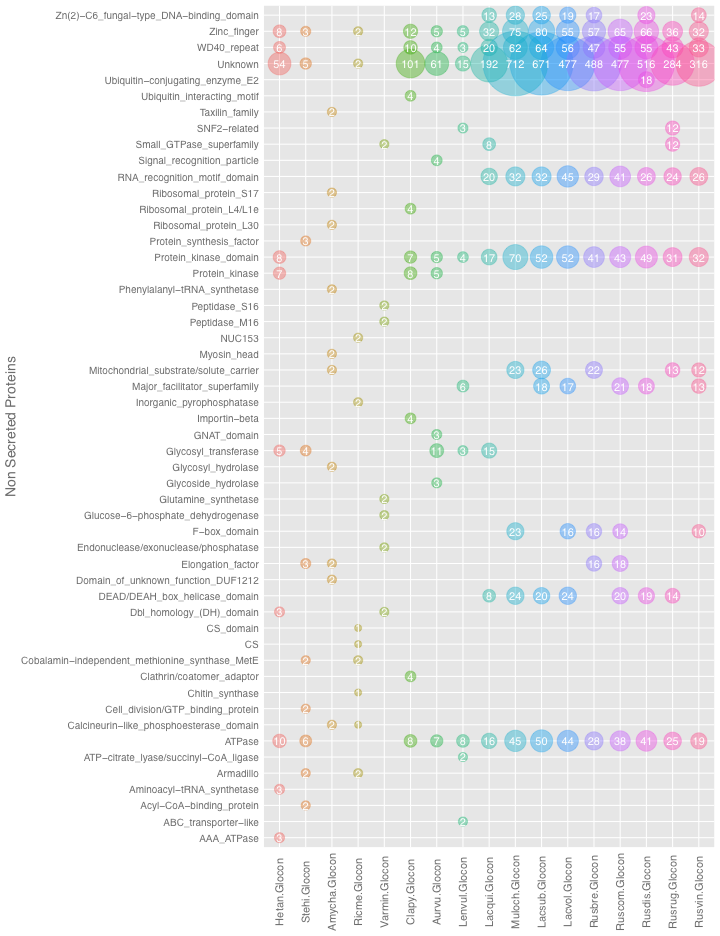

***Figure S12****.* ***Number of genes coding for non-secreted proteins in synteny****. Genes for non-secreted proteins per fungus in the syntenic regions were counted between Glocon and the rest of Russulales. Rusvin (R. vinacea), Rusrug (R. rugulosa), Rusbre (R. brevipes), Rusdis (R. dissimulans), Ruscom (R. compacta), Lacsub (L. subvellereus), Lacvol (L. volemus), Muloch (M. ochricompacta), Lacqui (L. quietus), Glocon (G. convolvens), Lenvul (L. vulpinus), Aurvu (A. vulgare), Clapy (C. pyxidata), Stehi (S. hirsutum), Hetan (H. annosum), Ricme (Peniophora sp.), Varmin (V. minispora), Amycha (A. chailletii).*

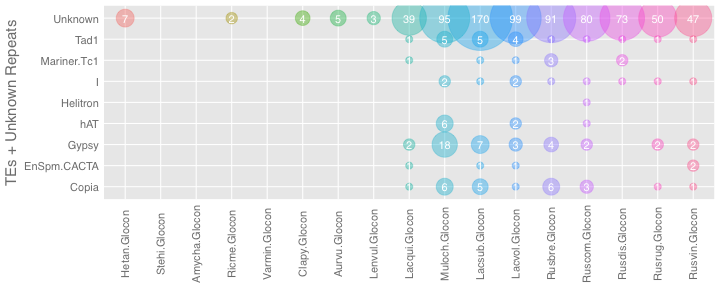

***Figure S13****.* ***Number of transposable elements in synteny****. The repeat elements were counted in the shared syntenic regions between Glocon and the rest of Russulales. Rusvin (R. vinacea), Rusrug (R. rugulosa), Rusbre (R. brevipes), Rusdis (R. dissimulans), Ruscom (R. compacta), Lacsub (L. subvellereus), Lacvol (L. volemus), Muloch (M. ochricompacta), Lacqui (L. quietus), Glocon (G. convolvens), Lenvul (L. vulpinus), Aurvu (A. vulgare), Clapy (C. pyxidata), Stehi (S. hirsutum), Hetan (H. annosum), Ricme (Peniophora sp.), Varmin (V. minispora), Amycha (A. chailletii).*

All identified classes of TEs only share synteny amongst Russulaceae and most unknown repeats are shared amongst Russulaceae. Shared syntenies with *H. irregulare* vs *G. convolvens* and *S. hirsutum* vs *G. convolvens* had no TEs, but there are only a few unknown repeats in *H irregulare* (Fig S13). The highest amount of TEs was seen when *G. convolvens* was compared to the closely related fungi including *M. ochricompacta*, *Lf. subvellereus*, *Lf. volemus*, *Russula* species.

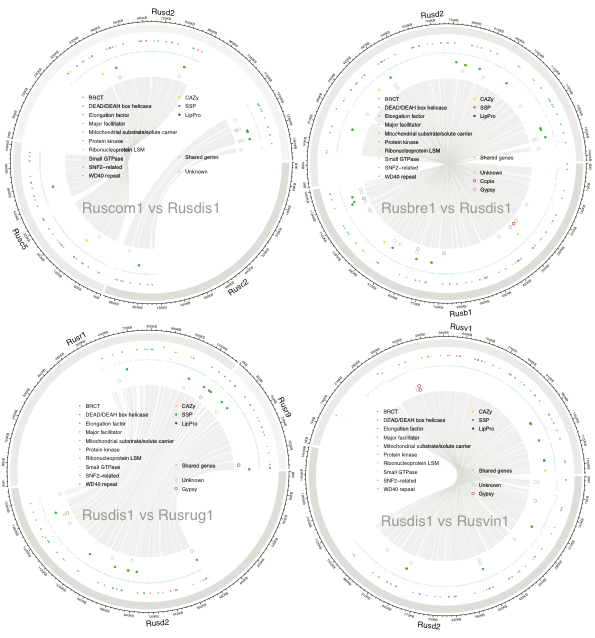

***Figure S14. Mesosynteny Hanabi plots with five Russula****. Rusvin1 (R. vinacea), Rusrug1 (R. rugulosa), Rusbre1 (R. brevipes), Rusdis1 (R. dissimulans), Ruscom1 (R. compacta).* ***Outer circle****: Rusdis scaffold 2 aligned with Russula scaffolds.* ***First inner circle****: Top 10 most frequent genes found among 5 Russula (see the legend).* ***Second inner circle****: The rest of genes shared among 5 Russula.* ***Third inner circle****: CAZymes, SSPs, lipases + proteases.* ***Fourth inner circle****: TEs and unknown repeats.* ***Links****: Mesosyntenic blocks based on similar sequences*

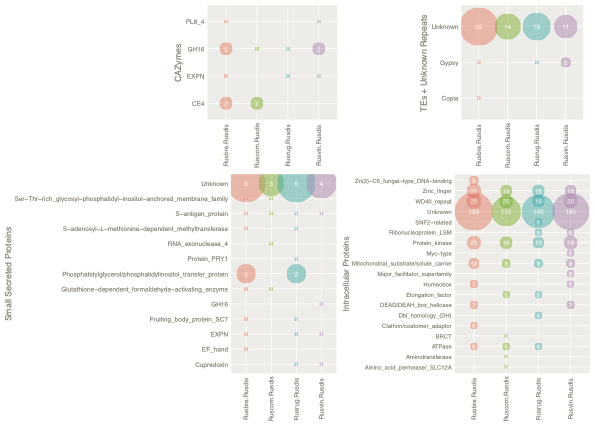

***Figure S15****.* ***Number of genes in Mesosynteny****. The count of the gene coding for CAZyme, SSPs, lipase+protease, intracellular proteins from the all microsyntenic blocks were indentified. Rusvin (R. vinacea), Rusrug (R. rugulosa), Rusbre (R. brevipes), Rusdis (R. dissimulans), Ruscom (R. compacta), Lacsub (L. subvellereus), Lacvol (L. volemus), Muloch (M. ochricompacta), Lacqui (L. quietus), Glocon (G. convolvens), Lenvul (L. vulpinus), Aurvu (A. vulgare), Clapy (C. pyxidata), Stehi (S. hirsutum), Hetan (H. annosum), Ricme (Peniophora sp.), Varmin (V. minispora), Amycha (A. chailletii).* ***NOTE****: Top 10 most frequent genes are used for intracellular proteins. Details of the individual count are given separately (Supplementary Data: 5Russula_Step14.3_MicroSynteny_CountSummary.csv).*

The orientation of a syntenic block is conserved in all but *R. rugulosa*, indicating a potential inversion for *R. rugulosa*. *Russula brevipes* and *R. dissimulans* share the highest level of mesosynteny containing a total of 367 shared gene annotations (Table S# - 5Russula_Step14.3_MicroSynteny_CountSummary.csv). Although a majority of syntenic genes are inferred as intracellular, some number of repeats, CAzymes, and SSPs are detected. In a large cluster of genes within this block, 22 genes with identical annotations are shared with only 3-4 orphan genes present in either genome (Fig. S15).

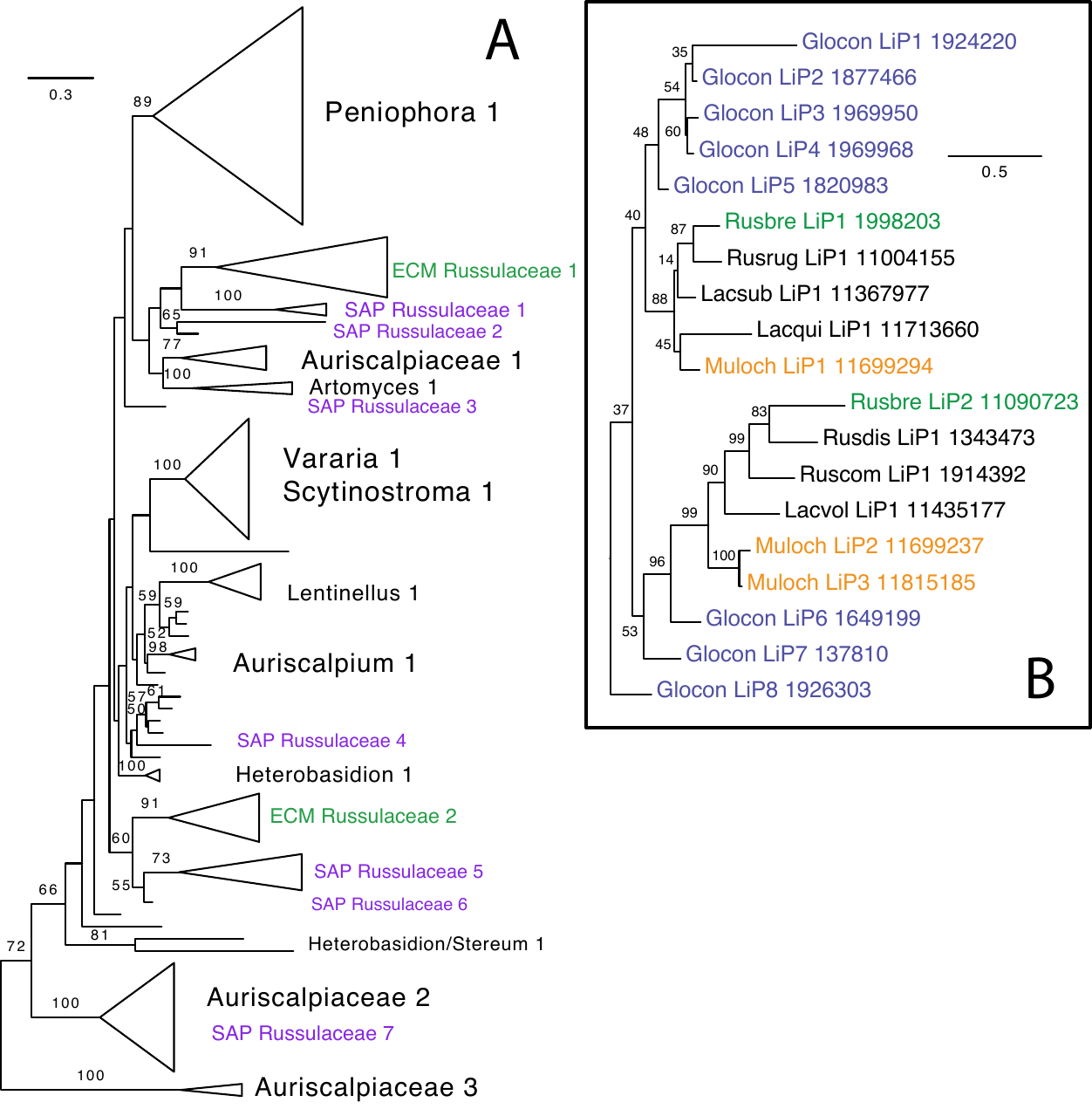

***Figure S16. Phylogenetic reconstruction of the class-II peroxidase (POD) gene family from Russulales that codes for lignin peroxidase. A)*** *Well-supported clades (bootstrap ≥70) are named based on included taxa. Clades including or made up of members of Russulaceae are colored based on nutritional mode (green–ectomycorrhizal; purple–saprotrophic).* ***B)*** *Phylogenetic reconstruction of the POD gene family only from Russulaceae. Species with multiple gene copies are highlighted in color, including two ectomycorrhizal species (*M. ochricompacta *&* R. brevipes*) and the saprotrophic representative (*G. convolvens*). Abbreviations for genomes are JGI identifiers (see Fig. S1).*

The gene history for the lignin peroxidases was inferred with 1000 bootstrap replicates. A phylogenetic reconstruction of class-II peroxidases from the Russulales resolved seven clades of saprotrophic Russulaceae (*G. convolvens*) and two clades of ECM Russulaceae with good support (bootstrap ≥70). All ECM members of Russulaceae are represented with at least one POD gene, with *Russula brevipes* and *Multifurca ochricompacta* having representative genes in each clade. It appears that *Multifurca ochricompacta* has recently undergone a gene duplication event.

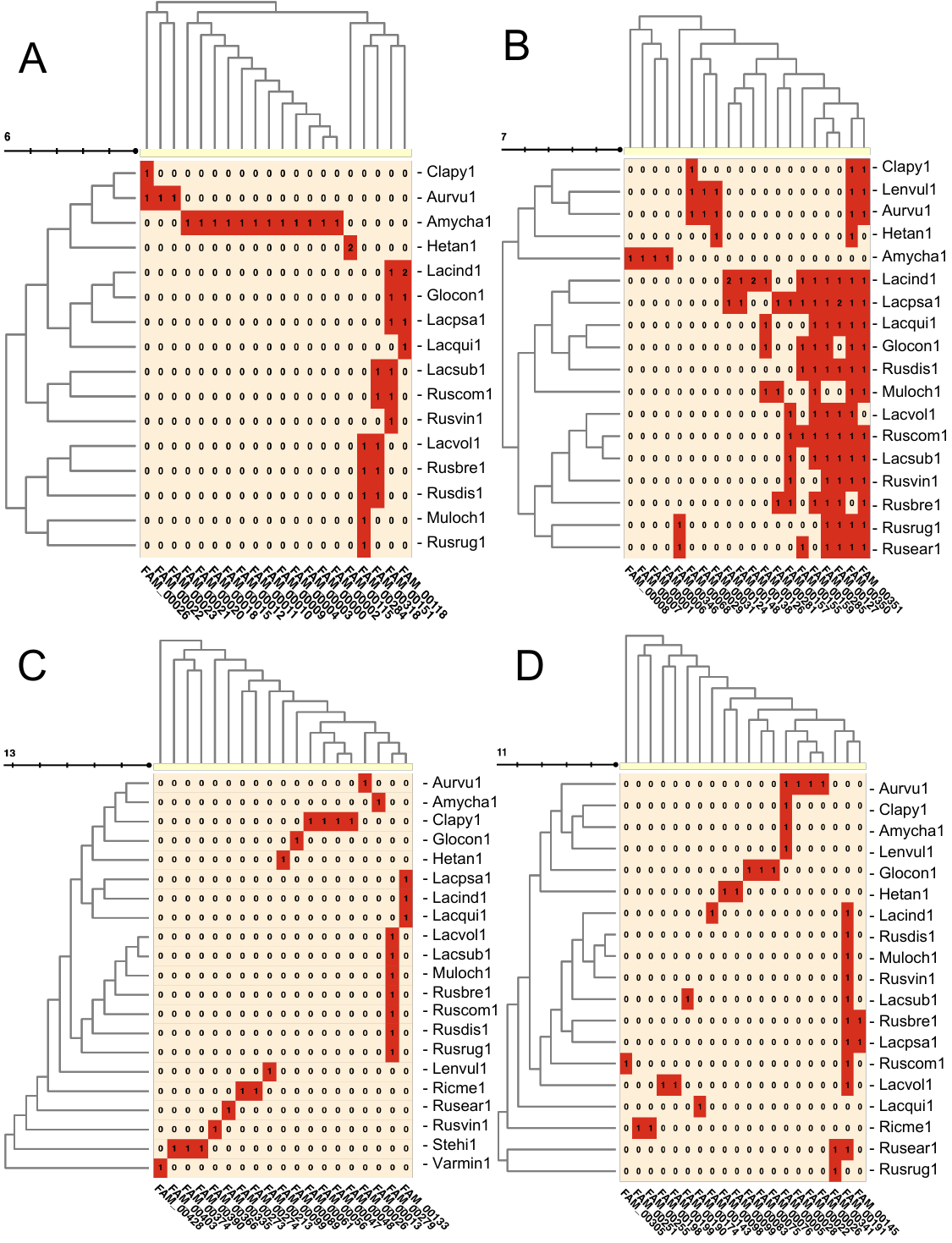

***Figure S17. Secondary metabolite cluster homology analysis for different classes of SMCs****.* ***A)*** *The top 20 most inclusive homologous groups of NRPS clusters.* ***B)*** *The top 20 most inclusive homologous groups of terpene clusters.* ***C)*** *The top 20 most inclusive homologous groups of PKSI clusters.* ***D)*** *The top 20 most inclusive homologous groups of other clusters. The number of clusters are given in red boxes and Pfam HMM cluster annotation is given below. Scale is given for number of node levels for species phylogeny based on presence/absence.*
