## Supplementary material for "Evolutionary priming and transition to the ectomycorrhizal habit in an iconic lineage of mushroom-forming fungi: is preadaptation a requirement?": Table S14

**Supplementary Table #**

| Fungal ID | Gene Type | P value | Random Mean Distance | Genome Mean Distance | Difference |
| --- | --- | --- | --- | --- | --- |
| Hetan | CAZymes | 0.373 | 19166 | 19797 | 632 |
| Stehi | CAZymes | 0.019 | 12983 | 15189 | 2206 |
| Amycha | CAZymes | 0.422 | 12414 | 12052 | -362 |
| Varmin | CAZymes | 0.455 | 8306 | 8168 | -138 |
| Ricme | CAZymes | 0.080 | 21021 | 23604 | 2583 |
| Clapy | CAZymes | 0.243 | 20893 | 22551 | 1658 |
| Aurvu | CAZymes | 0.392 | 11539 | 11845 | 306 |
| Lenvul | CAZymes | 0.189 | 11801 | 12932 | 1132 |
| Glocon | CAZymes | 0.525 | 15208 | 15200 | -8 |
| Lacqui | CAZymes | 0.109 | 4673 | 3181 | -1492 |
| Muloch | CAZymes | 0.091 | 18632 | 14124 | -4507 |
| Lacsub | CAZymes | 0.142 | 5586 | 4288 | -1297 |
| Lacvol | CAZymes | 0.174 | 4457 | 3541 | -917 |
| Rusbre | CAZymes | 0.119 | 5870 | 4557 | -1313 |
| Ruscom | CAZymes | 0.156 | 10550 | 8337 | -2214 |
| Rusdis | CAZymes | 0.438 | 7008 | 6655 | -353 |
| Rusrug | CAZymes | 0.159 | 5666 | 6859 | 1193 |
| Rusvin | CAZymes | 0.302 | 9758 | 8003 | -1755 |
| Hetan | Intracellular | 0.219 | 19193 | 19154 | -39 |
| Stehi | Intracellular | 0.087 | 12962 | 12919 | -43 |
| Amycha | Intracellular | 0.500 | 12415 | 12415 | 0 |
| Varmin | Intracellular | 0.010 | 8288 | 8341 | 52 |
| Ricme | Intracellular | 0.152 | 21067 | 21015 | -52 |
| Clapy | Intracellular | 0.232 | 20984 | 21026 | 42 |
| Aurvu | Intracellular | 0.268 | 11548 | 11565 | 17 |
| Lenvul | Intracellular | 0.453 | 11795 | 11799 | 4 |
| Glocon | Intracellular | 0.066 | 15262 | 15341 | 79 |
| Lacqui | Intracellular | 0.000 | 4683 | 4777 | 94 |
| Muloch | Intracellular | 0.000 | 18603 | 18923 | 321 |
| Lacsub | Intracellular | 0.000 | 5606 | 5674 | 68 |
| Lacvol | Intracellular | 0.000 | 4472 | 4541 | 69 |
| Rusbre | Intracellular | 0.000 | 5864 | 5922 | 58 |
| Ruscom | Intracellular | 0.000 | 10564 | 10693 | 129 |
| Rusdis | Intracellular | 0.002 | 7061 | 7112 | 51 |
| Rusrug | Intracellular | 0.000 | 5712 | 5808 | 96 |
| Rusvin | Intracellular | 0.000 | 9811 | 9957 | 146 |
| Hetan | Lipase/Protease | 0.010 | 19053 | 29457 | 10404 |
| Stehi | Lipase/Protease | 0.205 | 12954 | 14568 | 1614 |
| Amycha | Lipase/Protease | 0.047 | 12459 | 16839 | 4380 |
| Varmin | Lipase/Protease | 0.195 | 8297 | 7044 | -1253 |
| Ricme | Lipase/Protease | 0.438 | 21046 | 21342 | 296 |
| Clapy | Lipase/Protease | 0.157 | 20912 | 17186 | -3726 |
| Aurvu | Lipase/Protease | 0.091 | 11567 | 9345 | -2222 |
| Lenvul | Lipase/Protease | 0.403 | 11837 | 11203 | -634 |
| Glocon | Lipase/Protease | 0.103 | 15169 | 10710 | -4459 |
| Lacqui | Lipase/Protease | 0.000 | 4682 | 1588 | -3094 |
| Muloch | Lipase/Protease | 0.000 | 18674 | 6057 | -12616 |
| Lacsub | Lipase/Protease | 0.201 | 5622 | 4361 | -1261 |
| Lacvol | Lipase/Protease | 0.000 | 4480 | 1572 | -2908 |
| Rusbre | Lipase/Protease | 0.307 | 5887 | 4858 | -1030 |
| Ruscom | Lipase/Protease | 0.002 | 10600 | 3606 | -6994 |
| Rusdis | Lipase/Protease | 0.454 | 7085 | 6657 | -428 |
| Rusrug | Lipase/Protease | 0.003 | 5679 | 2262 | -3418 |
| Rusvin | Lipase/Protease | 0.408 | 9787 | 8594 | -1193 |
| Hetan | SSPs | 0.350 | 19143 | 18314 | -829 |
| Stehi | SSPs | 0.320 | 12956 | 12476 | -480 |
| Amycha | SSPs | 0.250 | 12409 | 11582 | -827 |
| Varmin | SSPs | 0.002 | 8293 | 6451 | -1842 |
| Ricme | SSPs | 0.396 | 21026 | 21380 | 354 |
| Clapy | SSPs | 0.191 | 20948 | 19279 | -1669 |
| Aurvu | SSPs | 0.383 | 11572 | 11248 | -324 |
| Lenvul | SSPs | 0.219 | 11800 | 10950 | -850 |
| Glocon | SSPs | 0.084 | 15248 | 13019 | -2229 |
| Lacqui | SSPs | 0.000 | 4689 | 2600 | -2089 |
| Muloch | SSPs | 0.000 | 18596 | 9152 | -9445 |
| Lacsub | SSPs | 0.000 | 5605 | 3135 | -2470 |
| Lacvol | SSPs | 0.000 | 4468 | 2392 | -2075 |
| Rusbre | SSPs | 0.000 | 5851 | 3941 | -1910 |
| Ruscom | SSPs | 0.000 | 10553 | 6222 | -4331 |
| Rusdis | SSPs | 0.001 | 7068 | 4704 | -2364 |
| Rusrug | SSPs | 0.000 | 5715 | 2669 | -3045 |
| Rusvin | SSPs | 0.000 | 9787 | 4588 | -5200 |

***Supplementary Table #****. Differences of TE-gene mean distances (= averaged distances in 5,000 random models subtracted from mean distances in actual genomes) for SSPs, CAZymes, lipases+proteases, and intracellular genes.* ***Orange****: Entries with statistical significance (p<0.01).*
