## Supplementary material for "Evolutionary priming and transition to the ectomycorrhizal habit in an iconic lineage of mushroom-forming fungi: is preadaptation a requirement?": Table S16

| CAFÉ # | # of Fungi | # of Copies | Annotation |
| --- | --- | --- | --- |
| **Russulaceae** | | | |
| 283 | 17 | 27 | GH5_15 β-1,6-glucanase |
| 286 | 8 | 27 | Aspartic_protease |
| 4772 | 5 | 16 | Subtilase_family |
| 5185 | 8 | 15 | GH27 α-galactosidase |
| 5345 | 9 | 14 | CBM1-GH5_7 β-mannanase |
| 5741 | 8 | 12 | GH3 β-glucosidase |
| 6265 | 8 | 10 | GH95 α-1,2-L-fucosidase |
| 6735 | 6 | 9 | Glutathione_S-transferase_N-terminal_domain |
| 6797 | 5 | 9 | AA3_2 aryl alcohol oxidase |
| 6802 | 8 | 9 | N-acetylglucosaminyl-phosphatidylinositol_de-N-acetylase |
| 6819 | 8 | 9 | GH76 endo-α-1,6-mannanase |
| 6953 | 8 | 8 | GH12endo-β-1,4-glucanase |
| 7230 | 8 | 8 | GH3 β-glucosidase |
| 7261 | 8 | 8 | GH145 α-L-rhamnohydrolase |
| 7282 | 8 | 8 | Putative_GTP_cyclohydrolase_1_type_2 |
| 7295 | 8 | 8 | GH43 endo-α-1,5-L-arabinosidase |
| 7310 | 7 | 8 | Aspartic_protease |
| 7673 | 6 | 7 | GH28 polygalacturonase |
| 7699 | 7 | 7 | CE8 pectin methylesterase |
| 7703 | 7 | 7 | EXPN |
| 7744 | 6 | 7 | Cerato-platanin |
| 7788 | 6 | 7 | GH53 endosialidase |
| 7832 | 5 | 7 | EXPN |
| 8217 | 5 | 6 | Subtilase_family |
| **ECM Russulaceae** | | | |
| 19 | 11 | 95 | Subtilase_family |
| 124 | 10 | 37 | AA3_2 aryl alcohol oxidase |
| 326 | 9 | 25 | GH7-CBM1 cellobiohydrolase |
| 581 | 9 | 21 | CBM1-GH10 xylanase |
| 4229 | 9 | 17 | GH55 exo-β-1,3-glucanase |
| 4849 | 9 | 16 | GH51 α-L-arabinofuranosidase |
| 5345 | 9 | 14 | CBM1-GH5_7 β-mannanase |
| 5400 | 8 | 14 | CBM5 chitinase |
| 5564 | 9 | 13 | GH2 β-glucuronidase |
| 5571 | 9 | 13 | AA8-AA3_1 cellobiose dehydrogenases |
| 5740 | 7 | 12 | GH18 chitinase |
| 5949 | 9 | 11 | Carboxylesterase_family |
| 5964 | 9 | 11 | GH12 endo-β-1,4-glucanase |
| 5974 | 9 | 11 | GH74-CBM1 xyloglucanase |
| 5980 | 9 | 11 | AA9-CBM1 lytic polysaccharide monooxygenase |
| 5986 | 9 | 11 | GH115 *a*-(4-O-methyl-)glucuronidase |
| 6202 | 9 | 10 | GH28 polygalacturonase |
| 6203 | 8 | 10 | CE8 pectin methylesterase |
| 6207 | 9 | 10 | GH131-CBM1 exo-β-1,3/1,6-glucanase |
| 6209 | 9 | 10 | CBM1-GH6 cellobiohydrolase |
| 6215 | 9 | 10 | GH71 α-1,3-glucanase |
| 6230 | 8 | 10 | CE16 acetyl esterase |
| 6235 | 9 | 10 | GH79 β-glucuronidase |
| 6237 | 8 | 10 | CBM1-CE16 acetyl esterase |
| 6244 | 9 | 10 | AA9 lytic polysaccharide monooxygenase |
| 6280 | 9 | 10 | GH18-CBM5 chitinase |
| 6743 | 9 | 9 | Fungal_cellulose_binding_domain |
| 6766 | 9 | 9 | Pyridoxamine_5 |
| 6777 | 9 | 9 | GH43-CBM35 exo-β-1,3-galactanase |
| 6779 | 9 | 9 | AA9-CBM1 lytic polysaccharide monooxygenase |
| 6793 | 9 | 9 | AA9-CBM1 lytic polysaccharide monooxygenase |
| 6826 | 9 | 9 | GH79 β-glucuronidase |
| 6867 | 6 | 9 | CFEM_domain |
| 7208 | 7 | 8 | GH128 endo-β-1,3-glucanase |
| 7228 | 8 | 8 | Serine_carboxypeptidase |
| 7297 | 8 | 8 | Carboxymuconolactone_decarboxylase_family |
| 7316 | 8 | 8 | Fungal_cellulose_binding_domain |
| 7340 | 5 | 8 | AA2 lignin peroxidase |
| 7365 | 6 | 8 | GH81 endo-β-1,3-glucanase |
| 7733 | 6 | 7 | AA9 lytic polysaccharide monooxygenase |
| 7736 | 7 | 7 | CE16 acetyl esterase |
| 8229 | 6 | 6 | Uncharacterized_oxidoreductase_C162.03 |
| 8274 | 6 | 6 | GH5_50 endo-β-1,4-glucanase |
| 8294 | 6 | 6 | AA9 lytic polysaccharide monooxygenase |
| **Russ/Lactif/Multi** | | | |
| 5943 | 10 | 11 | GH18 chitinase |
| 6276 | 10 | 10 | Altered_inheritance_of_mitochondria_protein_6 |
| **Russula/Lactifluus** | | | |
| 61 | 12 | 53 | AA3_2 aryl alcohol oxidase |
| 87 | 11 | 44 | Serine_carboxypeptidase_S28 |
| 321 | 10 | 25 | Extracellular_metalloproteinase |
| 5727 | 10 | 12 | Protein_of_unknown_function_(DUF1524) |
| 5752 | 10 | 12 | Phosphatidylethanolamine-binding_protein |
| 5783 | 9 | 12 | AA5_1 glyoxal oxidase |
| 5956 | 11 | 11 | AA3_2 aryl alcohol oxidase |
| 5987 | 10 | 11 | GH125 exo-α-1,6-mannosidase |
| 6205 | 10 | 10 | Probable_zinc_metalloprotease |
| 6264 | 10 | 10 | Domain_of_unknown_function_(DUF1932) |
| 6288 | 9 | 10 | Ser-Thr-rich_glycosyl-phosphatidyl-inositol-anchored_membrane_family |
| 6870 | 8 | 9 | Ferritin-like_domain |
| 7319 | 7 | 8 | PL14_4 β-1,4-glucuronan lyase |
| 7836 | 7 | 7 | Ser-Thr-rich_glycosyl-phosphatidyl-inositol-anchored_membrane_family |
| **Lactifluus** | | | |
| 5786 | 11 | 12 | Extracellular_metalloprotease |
| 5788 | 12 | 12 | GH18-CBM5 chitinase |
| 5813 | 8 | 12 | AA1_1 laccase |
| 6282 | 9 | 10 | PL14; PL14_3 β-1,4-glucuronan lyase |
| 6839 | 9 | 9 | Cerato-platanin |
| 6871 | 9 | 9 | EXPN |
| **Russula** | | | |
| 7 | 17 | 159 | Carboxylesterase_family |
| 242 | 13 | 29 | Eukaryotic_aspartyl_protease |
| 1681 | 5 | 19 | Serine_carboxypeptidase |
| 2008 | 10 | 18 | GH30_3 β-1,6-glucanase |
| 2137 | 12 | 18 | GH92 α-mannosidase |
| 5351 | 12 | 14 | GH92 α-mannosidase |
| 5363 | 11 | 14 | Thaumatin_family |
| 5553 | 13 | 13 | Serine_carboxypeptidase |
| 5743 | 12 | 12 | Ser-Thr-rich_glycosyl-phosphatidyl-inositol-anchored_membrane_family |
| **Russula exterior** | | | |
| 5472 | 14 | 14 | Fasciclin_domain |
| 5547 | 12 | 13 | Domain_of_unknown_function_(DUF4748) |
| 5773 | 12 | 12 | GH18 chitinase |
| 5813 | 8 | 12 | AA1_1 laccase |
| 5814 | 11 | 12 | Lipase_(class_3) |
| 6210 | 7 | 10 | CBM50x2-M23B |
| 6231 | 10 | 10 | Ser-Thr-rich_glycosyl-phosphatidyl-inositol-anchored_membrane_family |
| 6839 | 9 | 9 | Cerato-platanin |
| 6865 | 9 | 9 | Protein_priA |
| **Russula interior** | | | |
| 4978 | 14 | 16 | CBM1-GH45 endoglucanase |
| 5554 | 7 | 13 | Carboxylesterase_family |
| 5786 | 11 | 12 | Extracellular_metalloprotease |
| 6208 | 10 | 10 | EXPN |
| 6282 | 9 | 10 | PL14_3 β-1,4-glucuronan lyase |
| 6871 | 9 | 9 | EXPN |

***Table S#. Orthologous gene clusters contracted in the ancestry of Russulales.***

We detect a loss of carbohydrate binding modules (CBMs), glycoside hydrolases (GHs), and polysaccharide lyases (PLs) for ECM Russulaceae. More specifically the ancestor of the ECM Russulaceae experienced a significant contraction in 44 gene clusters, including multiple clusters of AA2 lignin peroxidases, AA9 (formerly GH61) lytic polysaccharide mono-oxygenases, GH5_5 endoglucanases, CBM1 cellulose binding modules, expansins, and a diverse array of GH glycoside hydrolase families. This contracted loss of secreted enzymes is also seen in the ancestor of Russulaceae, with 24 significant contractions that includes groups of AA3_2 aryl alcohol oxidases, CE8 pectin methylesterases, many glycoside hydrolase families, expansins, and aspartyl proteases. Another large contraction of secreted proteins is seen in the ancestor of *Russula* and *Lactifluus*, with 14 significant gene cluster contractions, including multiple clusters of AA and GH groups, subtilases, and P14_4 glucuronan lyases.
