## Supplementary material for "Evolutionary priming and transition to the ectomycorrhizal habit in an iconic lineage of mushroom-forming fungi: is preadaptation a requirement?": S17

| CAFÉ # | # of Fungi | # of Copies | Annotation |
| --- | --- | --- | --- |
| **Russulaceae** | | | |
| 5674 | 9 | 13 | AA1 1 laccase |
| 5692 | 10 | 13 | Eukaryotic aspartyl protease |
| 6070 | 9 | 11 | Double-stranded RNA binding motif |
| 6099 | 10 | 11 | MYND finger |
| 6267 | 10 | 10 | GH16; Probable glycosidase C21B10.07 |
| 6320 | 7 | 10 | AA2 lignin peroxidase |
| 6367 | 9 | 10 | GH5 9 exo-β-1,3-glucanase |
| 6416 | 10 | 10 | CE14; N-acetylglucosaminyl-phosphatidylinositol de-N-acetylase |
| 6440 | 8 | 10 | Uncharacterized protein AFUA 6G02800 |
| 6515 | 8 | 10 | Lipase (class 3) |
| 6619 | 10 | 10 | Domain of unknown function (DUF3844) |
| 6633 | 10 | 10 | RNA recognition motif. (a.k.a. RRM RBD or RNP domain) |
| 7396 | 7 | 8 | Protein of unknown function (DUF3445) |
| 7445 | 8 | 8 | CFEM domain |
| 7447 | 8 | 8 | Peptidase family M28; Peptidase family M20/M25/M40 |
| 7883 | 7 | 7 | Ser-Thr-rich glycosyl-phosphatidyl-inositol-anchored membrane family |
| **ECM Russulaceae** | | | |
| 83 | 18 | 45 | AA5 1 glyoxal oxidase |
| 202 | 17 | 31 | GH18 chitinases |
| 242 | 13 | 29 | Eukaryotic aspartyl protease |
| 433 | 15 | 23 | GH47 α-mannosidases |
| 1681 | 5 | 19 | Serine carboxypeptidase |
| 5931 | 11 | 11 | Ser-Thr-rich glycosyl-phosphatidyl-inositol-anchored membrane family |
| 6747 | 9 | 9 | CFEM domain |
| 7055 | 8 | 9 | EXPN |
| 7488 | 7 | 8 | PL14 β-1,4-glucuronan lyase |
| **Russula/Lactifluus/Multifurca** | | | |
| 8707 | 5 | 6 | Lipase (class 3) |
| **Lactifluus** | | | |
| 344 | 18 | 25 | AA1 2 ferroxidase |
| 968 | 17 | 19 | Phosphatidylethanolamine-binding protein |
| 1211 | 17 | 19 | AA2 lignin peroxidase |
| **Russula** | | | |
| 735 | 16 | 20 | Peroxidase family 2 |
| 7622 | 4 | 8 | Aspartic protease |
| **Russula exterior** | | | |
| 344 | 18 | 25 | AA1 2 ferroxidase |
| **Russula interior** | | | |
| 191 | 16 | 32 | Serine carboxypeptidase |
| 336 | 12 | 25 | Carboxylesterase family |
| 400 | 15 | 23 | Carboxylesterase family |

***Table S#. Orthologous gene clusters expanded in the ancestry of Russulales.***

Secretome specialization for ECM Russulaceae is characterized by significant expansions in gene copy number for orthologous clusters (= orthogroups) of aspartyl proteases, subtilisin-like serine proteases, serine carboxypeptidases, AA5_1 glyoxyl oxidases, GH47 α-mannosidases, GH18 chitinases, and others for a total of 9 expanded orthologous clusters (Table S#). The ancestor of Russulaceae saw an expansion in clusters of AA1_1 laccases, aspartyl proteases, AA2 lignin peroxidases, GH5_9 exo-β-1,3-glucanases, class 3 lipases, and others for a total of 16 expanded orthologous clusters. One orthologous cluster, a group of class 3 lipases, was expanded in the ancestor of ECM Russulaceae excluding *Lactarius*. No expansions and many contractions were found for the shared ancestor of *Russula* and *Lactifluus*. Three orthologous clusters were found expanded in the ancestor of *Lactifluus*, including a group of AA1_2 ferroxidases and AA2 lignin peroxidases. The ancestor of *Russula* saw an expansion in a group of aspartic proteases and peroxidases.
