## Appendix 1 for "Evolutionary priming and transition to the ectomycorrhizal habit in an iconic lineage of mushroom-forming fungi: is preadaptation a requirement?"

**Genome sequencing and assembly**

All genomes, except for those of 4 species described below, were sequenced using Pacific Biosciences (PacBio). Unamplified libraries were generated using PacBio standard template preparation protocol for creating >= 10kb libraries. 1-10 µg of gDNA was used to generate each library and the DNA was sheared using Covaris g-Tubes^TM^ to generate sheared fragments of >= 10kb in length or using Megarupter to generate sheared fragments of 30kb in length. The sheared DNA fragments were then prepared using Pacific Biosciences SMRTbell template preparation kit, where the fragments were treated with DNA damage repair, had their ends repaired so that they were blunt-ended, and 5' phosphorylated. Pacific Biosciences hairpin adapters were then ligated to the fragments to create the SMRTbell template for sequencing. The SMRTbell templates were then purified using exonuclease treatments and size-selected using AMPure PB beads or BluePippin (Sage Science). Sequencing primer was then annealed to the SMRTbell templates and Version P6 sequencing polymerase was bound to them. The prepared SMRTbell template libraries were then sequenced on a Pacific Biosciences RSII sequencer using Version C4 chemistry and 2 hour sequencing movie run times. One of the 2 *Lactarius psammicola* libraries, as well as the *Lactarius indigo* and *Russula earlei* libraries, were sequenced differently, using v3 sequencing primer, 1M v2 SMRT cells, and Version 2.0 sequencing chemistry with 6 & 10 hour sequencing movie run times. Whatever the library preparation or sequencing methods, the resulting gDNA reads were then assembled using FALCON version 0.0.2, 0.4.2, 0.7.3, or 1.8.8 (Chin 2016).

The *Amylostereum*, *Auriscalpium*, and *Vararia* genomes were sequenced using Illumina. 100 ng of gDNA was sheared to 300 bp fragments using the Covaris LE220 and size selected using SPRI beads (Beckman Coulter). The fragments were treated with end-repair, A-tailing, and ligation of Illumina compatible adapters (IDT, Inc) using the KAPA-Illumina library creation kit (KAPA biosystems). The prepared libraries were quantified using KAPA Biosystem’s next-generation sequencing library qPCR kit and run on a Roche LightCycler 480 real-time PCR instrument. The quantified libraries were then multiplexed with other libraries, and the pool of libraries was then prepared for sequencing on the Illumina HiSeq sequencing platform utilizing a TruSeq paired-end cluster kit v4, and Illumina’s cBot instrument to generate a clustered flow cell for sequencing. Sequencing of the flow cell was performed on the Illumina HiSeq2500 sequencer using HiSeq TruSeq SBS sequencing kits, v4, following a 2x150bp indexed run recipe. Except for *Artomyces* described below, the resulting illumina gDNA reads were then assembled together with Velvet (Zerbino and Birney 2008). The resulting assembly was used to simulate a long mate-pair library with insert 3000 ± 300 bp, which was then assembled together with the original Illumina library with ALLPATHS-LG version R42328, R46652, or R49403 (Gnerre et al 2011).

For *Artomyces*, in addition to the Regular Fragment Illumina library described above, an additional 4kb CLRS Illumina Regular Long Mate-Pair library was created. 5-6 µg of DNA was sheared using the Covaris g-TUBE™ and gel size selected for 4kb. The sheared DNA was treated with end repair and ligated with biotinylated adapters containing loxP. The adapter ligated DNA fragments were circularized via recombination by a Cre excision reaction (NEB). The circularized DNA templates were then randomly sheared using the Covaris LE220.The sheared fragments were treated with end repair and A-tailing using the KAPA-Illumina library creation kit (KAPA biosystems) followed by immobilization of mate pair fragments on streptavidin beads (Invitrogen). Illumina compatible adapters (IDT, Inc) were ligated to the mate pair fragments and 10 cycles of PCR was used to enrich for the final library (KAPA Biosystems). The libraries were then treated as with the above Regular Fragment libraries, except for following a 2x100bp indexed run recipe. The resulting Illumina gDNA reads were then assembled together with ALLPATHS-LG version R47710.

**Transcriptome sequencing and assembly**

Sequenced transcriptomes were used to assess the completeness of genome assemblies and to seed and assess all genome annotations except for that of *Vararia*, from which no RNA was available. All transcriptomes were sequenced using Illumina RNA-Seq with polyA selection.

Except for 5 species described below, stranded cDNA libraries were generated using the Illumina Truseq Stranded mRNA Library Prep kit. mRNA was purified from 100 ng or 1 µg of total RNA using magnetic beads containing poly-T oligos. mRNA was fragmented using divalent cations and high temperature. The fragmented RNA was reversed transcribed using random hexamers and SSII (Invitrogen) followed by second strand synthesis. The fragmented cDNA was treated with end-pair, A-tailing, adapter ligation, and 10 cycles of PCR. For *Lactarius indigo*, *Lactifluus* *subvellereus*, *Multifurca*, *Russula* *earlei*, and *Russula* *vinacea*, plate-based RNA sample prep was performed on the PerkinElmer Sciclone NGS robotic liquid handling system using Illumina TruSeq Stranded mRNA HT sample prep kit utilizing poly-A selection of mRNA following the protocol outlined by Illumina in their user guide and with the following conditions: total RNA starting material was 1 µg per sample and 8 cycles of PCR was used for library amplification.

Except for 1 species described below, the prepared transcriptome libraries were quantified using KAPA Biosystem’s next-generation sequencing library qPCR kit and run on a Roche LightCycler 480 real-time PCR instrument. The quantified libraries were then multiplexed with other libraries, and the pool of libraries was then prepared for sequencing on the Illumina HiSeq sequencing platform utilizing a TruSeq paired-end cluster kit v4, and Illumina’s cBot instrument to generate a clustered flow cell for sequencing. Sequencing of the flow cell was performed on the Illumina HiSeq2500 sequencer using HiSeq TruSeq SBS sequencing kits v4, following a 2x150bp indexed run recipe. *Lactarius indigo* sequencing was performed on the Illumina NovaSeq sequencer using NovaSeq XP V1 reagent kits, a S4 flowcell, and a 2x150bp indexed run recipe. The resulting reads were assembled into RNA contigs using Trinity version 2.1.1 or 2.3.2 (Grabherr et al 2011), with the exception of *Artomyces* and *Auriscalpium*, whose RNA contigs were assembled using Rnnotator version 3.3.1 (Martin et al 2010).

**Genome Annotation**

Each genome was annotated using the JGI Annotation Pipeline, which detects and masks repeats and transposable elements, predicts genes, characterizes each conceptually translated protein with sub-elements such as domains and signal peptides, chooses a best gene model at each locus to provide a filtered working gene set, clusters the filtered sets into draft gene families, ascribes functional descriptions (such as GO terms and EC numbers), and creates a JGI genome portal in MycoCosm (https://mycocosm.jgi.doe.gov/) with tools for public access and community-driven curation of the annotation (Grigoriev *et al*., 2014; Kuo *et al*., 2014).
